# Cancer cells subvert the primate-specific KRAB zinc finger protein ZNF93 to control APOBEC3B

**DOI:** 10.1101/2025.03.05.641617

**Authors:** Romain Forey, Cyril Pulver, Charlène Raclot, Olga Rosspopoff, Sandra Offner, Julien Duc, Evarist Planet, Filipe Martins, Priscilla Turelli, Didier Trono

**Affiliations:** School of Life Sciences, École Polytechnique Fédérale de Lausanne, Lausanne, Switzerland; Nexco Analytics, Bâtiment Alanine, StarLab, Route de la Corniche 5A, 1066 Epalinges Switzerland

## Abstract

ZNF93 is a primate-restricted KRAB zinc finger protein responsible for repressing 20- to 12-million-year-old L1 transposable elements. Here, we reveal that ZNF93 also regulates the key cancer driver APOBEC3B—a mutagenic enzyme linked to tumorigenesis and cancer progression. ZNF93 depletion impairs DNA synthesis, activates replication and DNA damage checkpoints, and triggers proinflammatory phenotypes. Conversely, its overexpression enhances resistance to exogenous genotoxic stress, mirroring the effects observed with APOBEC3B depletion. ZNF93 expression correlates with cell proliferation rates and is overexpressed in many cancer types. These findings suggest that ZNF93 serves as a critical guardian of genome integrity, co-opted by cancer cells to counterbalance APOBEC3B-induced and L1-derived genomic instability and inflammation.

## INTRODUCTION

The human genome harbors >5 million inserts derived from transposable elements (TEs), which together contribute at least 60% of its DNA content (1). Amongst them are more than 1.5 million Long Interspersed Nuclear Element (LINE) integrants belonging in their majority to the LINE-1 (L1) subfamily (2). L1 are autonomous retrotransposons encoding the machinery necessary for their transposition, a “copy-and-paste” process whereby an RNA intermediate is reverse transcribed into a DNA copy that integrates back into the genome (3). Integration relies on the endonuclease activity of the L1-encoded ORF2p protein, contained in its N-terminal region, inducing DNA double-strand breaks as a first step (4). The C-terminal part of ORF2p then mediates reverse transcription, using the freed 3’ end of a DNA strand as a primer and the L1 RNA as a template (3).

L1 predate the origin of vertebrates and in mammals display a ladder-like phylogeny where only the youngest group is active at any given time, expanding from a common progenitor for a few million years before being replaced by a new group (2, 5). This pattern is consistent with an arms race between a host that evolves functions repressing L1 transposition, and L1 that mutates in return to bypass this obstacle (6). In the human lineage, L1 restriction factors are exemplified by ZNF93, a member of the Krüppel-associated box (KRAB)-containing zinc finger proteins (KZFP) family, the largest group of DNA-binding proteins in higher vertebrates (7, 8). There are close to 400 human KZFPs, the great majority of which recognize a particular TE subgroup as primary genomic target via a C-terminal array of zinc fingers, allowing for sequence-specific DNA binding (7–10). 80% of human KZFPs act as transcriptional repressors through recruitment by their N-terminal KRAB domain of a KAP1/TRIM28-nucleated heterochromatin-inducing complex while the remaining, often older family members display other functionalities linked to the presence of variant KRAB or additional domains (7, 8, 10, 11). ZNF93 is a canonical KZFP that emerged some 20 million years ago (mya), contemporary to the L1PA6 L1 subfamily, which it represses by recognizing a sequence located close to the 5’ end of their integrants (6). The ZNF93 target motif is conserved in the more recent L1PA5 and L1PA4 integrants but was lost through a 129bp-long deletion that occurred in some of their L1PA3 descendants, allowing these and their L1PA2 and L1Hs progeny to escape ZNF93-mediated control (6). While the ZNF93-L1 interplay is a convincing illustration of the arms race model, it poses a conundrum. Indeed, only ∼100 human L1 integrants are still replication-proficient, all belonging to the less than 3 million year old (myo) L1Hs subgroup (12). The TE targets of ZNF93, like those of an overwhelming majority of KZFPs, are thus transposition-incompetent, due to the accumulation of inactivating mutations over time, yet they remain bound by their cognate KZFP (6).

Cancer represents an interesting setting to study the interplay between TEs and their epigenetic controllers. In cancer cells, epigenetic alterations typically trigger the de-repression of TEs, which leads to DNA damage and secondary inflammation (13, 14). Cell clones are then selected that express high levels of epigenetic repressors, which re-instate TE silencing thereby mitigating replication stress and allowing cells to resume growth (15, 16). L1 have been identified as proinflammatory in the context of both cancer and cellular senescence (17, 18), whether due to loss of epigenetic repression in the former (16, 18), or to the effect of transcriptional activators in the latter (17). In both situations, increases in L1 expression correlate with elevated DNA damage and replication stress (16–18), part of which may be due to the endonuclease activity of ORF2p (4, 19, 20).

Here, we examined how L1-targeting KZFPs might influence this process. This led us to identify ZNF93 as a family member almost systematically upregulated in cancer cells. We further determined that many of the L1 targets of this KZFP encode a truncated form of ORF2p with increased endonuclease activity. Accordingly, we observed that ZNF93 depletion led to genomic instability and inflammation in cancer cells. However, we unexpectedly found that much of this effect stemmed from the loss of ZNF93-mediated repression of APOBEC3B, a cytidine deaminase known not only as a restriction factor of L1 (21–24) and various viruses (25), but also as an inducer of cell death upon DNA damage and replication stress (26–33) and a well-documented mutagen in cancer (31, 34–39). These results position ZNF93 as a critical guardian of genome stability subverted by cancer cells to foster tumor growth.

## RESULT

### KZFPs accumulate at the promoter of endonuclease-proficient young LINE-1 integrants

To identify KZFPs responsible for preventing L1 ORF2p-induced genotoxicity, we first conducted a census of KZFP binding over endonuclease-encoding (EN+) L1 elements. Most L1-derived sequences are 5’-truncated, thereby lacking the 5’ UTR-embedded promoter hence the ability to express L1-encoded proteins. We thus mapped KZFP binding over EN+ full length L1s, defined as L1 integrants spanning at least 5.9 kb and harboring an ORF2p open reading frame longer than 230 amino acids, the minimal length of its endonuclease moiety. We focused on the seven youngest L1 (YL1) subgroups, L1PA7 to L1HS (2), which together accounted for most of the full length L1s (6691/8597, 78%). We further determined that due the accumulation of inactivating mutations, only 28% (1842/6691) of full length YL1 (FLYL1) are EN+ (Fig. 1A).

**Figure 1:**
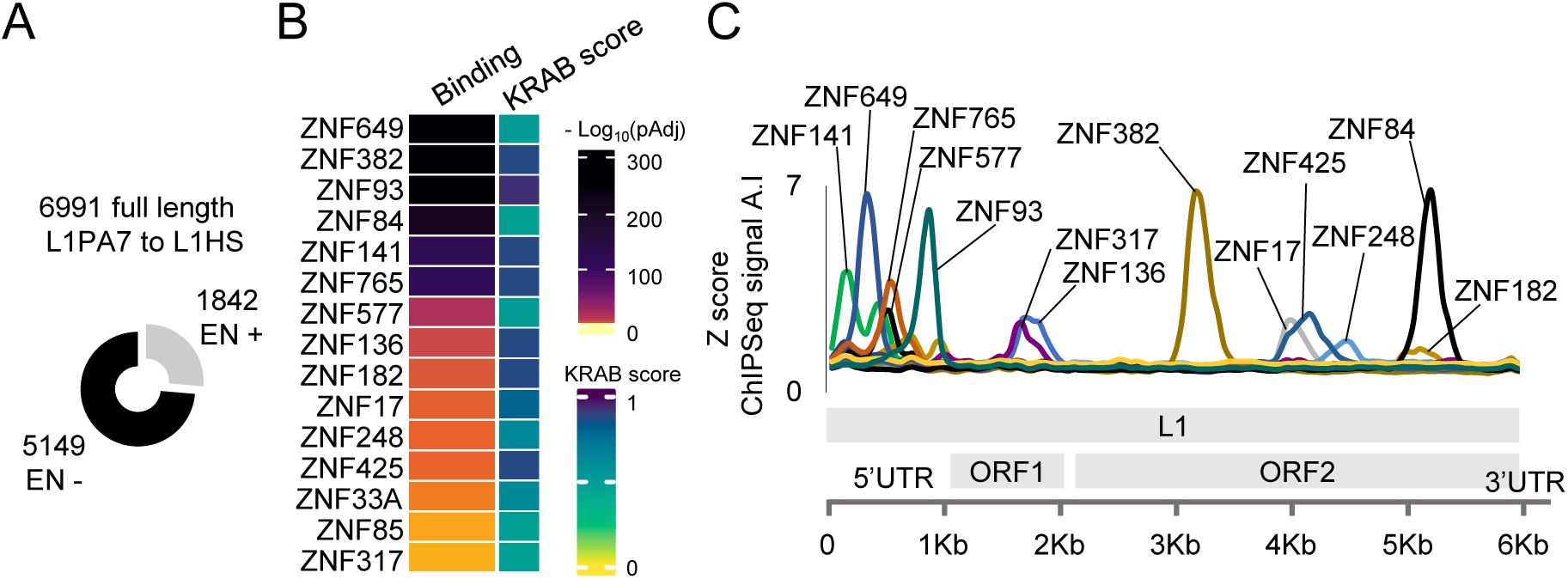
KZFPs accumulate at the promoter of endonuclease-proficient young LINE-1 integrants. A. Full-length (>5900 bp) and recently evolved (L1PA7 to L1HS) LINE-1 elements (FLYL1s), with (EN+) or without (EN−) putative endonuclease activity. B. Krüppel-associated box (KRAB)-containing zinc finger proteins (KZFP) binding enrichment (Fischer’s exact test, adj. p < 0.05) at FLYL1 elements endowed with endonuclease activity (FLYL1^EN+^). The KRAB score represents the probability of recruiting TRIM28 (Pulver, Forey *et al.*, 2025). Data from Imbeault *et al.*, 2017; Helleboid *et al.*, 2019; de Tribolet *et al.*, 2023. C. ChIP signal for the KZFPs identified in (B) across the meta-FLYL1^EN+^ sequence

Using previously compiled genome-wide binding profiles (7, 9, 10) we identified 15 KZFPs, the binding of which binding was enriched at FLYL1^EN+^ loci (Fig. 1B, S1A). All bore KRAB domains predicted to induce TRIM28-dependent H3K9me3 deposition (40), consistent with their potential role as FLYL1^EN+^ repressors. KZFP binding was particularly dense at FLYL1^EN+^ promoters, with ZNF649, ZNF93, ZNF765, ZNF577 and ZNF141 exhibiting the highest enrichments (Fig. 1C). Interestingly, KZFPs displayed preferential binding to specific regions of the FLYL1^EN+^ promoter. For example, ZNF93 predominantly targeted the 3’ end of the promoter, while ZNF765 localized significantly upstream in the 5’UTR and ZNF765 somewhere between them. Some KZFPs recognize regions outside of the promoter, such as ZNF382 and ZNF84, which bind to distant sequences in ORF2 (Fig. 1C).

Moreover, the KZFP-mediated recognition of FLYL1^EN+^ displayed remarkable subfamily-specificity (Fig. S1A). For instance, ZNF765 and ZNF93 showed maximal enrichment at L1PA5^EN+^ and L1PA4^EN+^ promoters, respectively, while ZNF382 and ZNF84 bound most FLYL1^EN+^ including from the L1HS subset. Also, L1PA7^EN+^ and L1HS^EN+^ - representing the oldest and youngest subfamilies of FLYL1, respectively – were recognized by the narrowest array of KZFPs (Fig. S1A). This pattern likely reflects on the one hand the progressive degeneration of KZFP-binding motifs in older subfamilies like L1PA7, and on the other hand insufficient evolutionary time for the selection and fixation of L1HS-specific KZFP repressors. Together, these results pointed to ZNF93, ZNF649 and ZNF765 as strong candidate FLYL1^EN+^ repressors.

### ZNF93 is expressed in proliferating cells, marking TRIM28 recruitment at FLYL1^EN+^ promoters

Increased expression of epigenetic repressors is frequently selected for in cancer cells, resulting in reduced TE expression, prevention of DNA damage, replication stress and inflammation, all of which partake in sustaining proliferation (15, 16, 18). Consequently, we reasoned that among FLYL1^EN+^-enriched KZFPs, those expressed concomitantly with proliferation would likely be the most functionally significant FLYL1^EN+^ repressors for cancer evolution.

To test this hypothesis, we evaluated the correlation between a 167-gene proliferation signature and the expression of FLYL1^EN+^-enriched KZFPs across 23 cancer subtypes from The Cancer Genome Atlas (TCGA). ZNF93 displayed the most pronounced (Wilcoxon two-tailed test, p < 0.05) and frequent (14/23 cancer subtypes) statistically significant (BH-adj. p < 0.05, t-test) association with proliferation (Fig. 2A). Notably, ZNF93 was only topped by ZNF695 and ZNF724P when considering expression-versus-proliferation correlations across all KZFPs and TCGA tumour samples (Fig. S2A). Moreover, ZNF93 displayed a greater tendency for upregulation in cancer versus normal tissue compared to other FLYL1^EN+^ promoter-enriched KZFPs (Fig. 2A). Together, these results shows that ZNF93 expression is correlated with proliferation in cancer.

**Figure 2:**
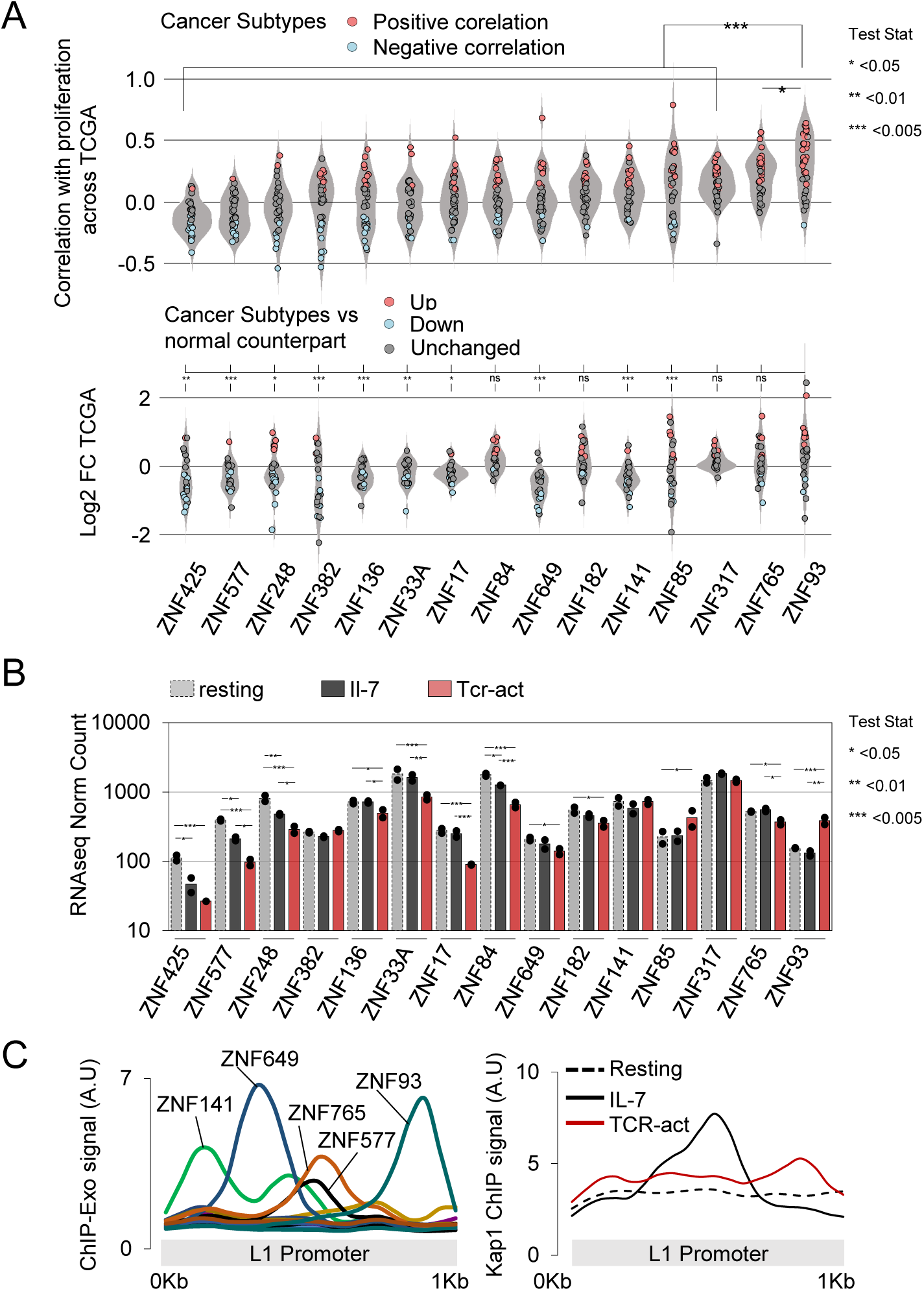
ZNF93 is expressed in proliferating cells, marking TRIM28 recruitment at FLYL1^EN+^ promoters. A. Top: Spearman rank correlation (t-test, adj. p < 0.05) between *KZFP* expression and a 167-gene proliferation signature across 33 TCGA cancer subtypes. Red: significant correlation; Blue: significant anticorrelation. Bottom: differential *KZFP* expression between cancer and normal samples (23 cancer subtypes selected). Median Spearman correlation coefficients and log2FC were compared across subtypes using two-tailed t-test. B. *KZFP* expression in resting, IL-7 activated, and TCR-activated CD4^+^T cells. Means across two independent experiments are shown. Differential expression analysis was performed. C. Left: ChIP signal for KZFPs across the meta-FLYL1^EN+^ promoter sequence. Right: TRIM28 ChIP signal in resting, IL-7 activated and TCR-activated CD4^+^T cells.

To explore further the link between FLYL1^EN+^-enriched KZFPs and proliferation, we analysed the differential responsiveness of resting primary CD4^+^T cells to IL-7, a non-mitogenic T cell activator, versus CD3/CD28 crosslinking, a strongly mitogenic T cell activator (public dataset (41), Fig. S2B). Genes upregulated by TCR crosslinking, but not IL-7, were enriched for cell cycle and proliferation markers (Fig. S2C, D). While IL-7 had little to no effect on the expression of FLYL1^EN+^-enriched KZFPs, most (13/15) were either unaffected or downregulated upon TCR crosslinking. In contrast ZNF93 was robustly upregulated upon TCR crosslinking (Fig. 2B). This result reinforces the association between ZNF93 expression and proliferation.

Next, we investigated whether binding of the KZFP cofactor TRIM28 to FLYL1^EN+^ loci was altered upon mitogenic versus non-mitogenic T cell activation. Strikingly, TRIM28 occupancy specifically increased at ZNF93 binding sites upon TCR crosslinking but not IL-7 treatment. In contrast, IL-7 treatment increased TRIM28 binding at ZNF649 and ZNF765 binding sites (Fig. 2C, S2E). These data indicate that among FLYL1EN+-enriched KZFPs, ZNF93 most specifically correlates with TRIM28 recruitment at L1 promoter under mitotic stimulation.

### ZNF93 depletion reduces proliferation and induces replicative stress in cancer cells

To assess the role of ZNF93 in proliferating cells, we depleted this KZFP through RNA interference, achieving a four-fold knockdown (KD) (Fig. 3A) in acute myeloid leukemia-derived K562 cells. *ZNF93* KD reduced the proliferation rate of K562 by ∼40% (Fig. 3B). This phenotype was consistently reproduced across multiple cell lines, including another leukemia cell line (THP1), three B-cell lymphoma cell lines (OCY19, SUDHL4 and U2932) and two colorectal cancer cell lines (LS1034 and SW480), each expressing various levels of the endogenous ZNF93 at baseline (Fig. 3C, D, S3A, B).

**Figure 3:**
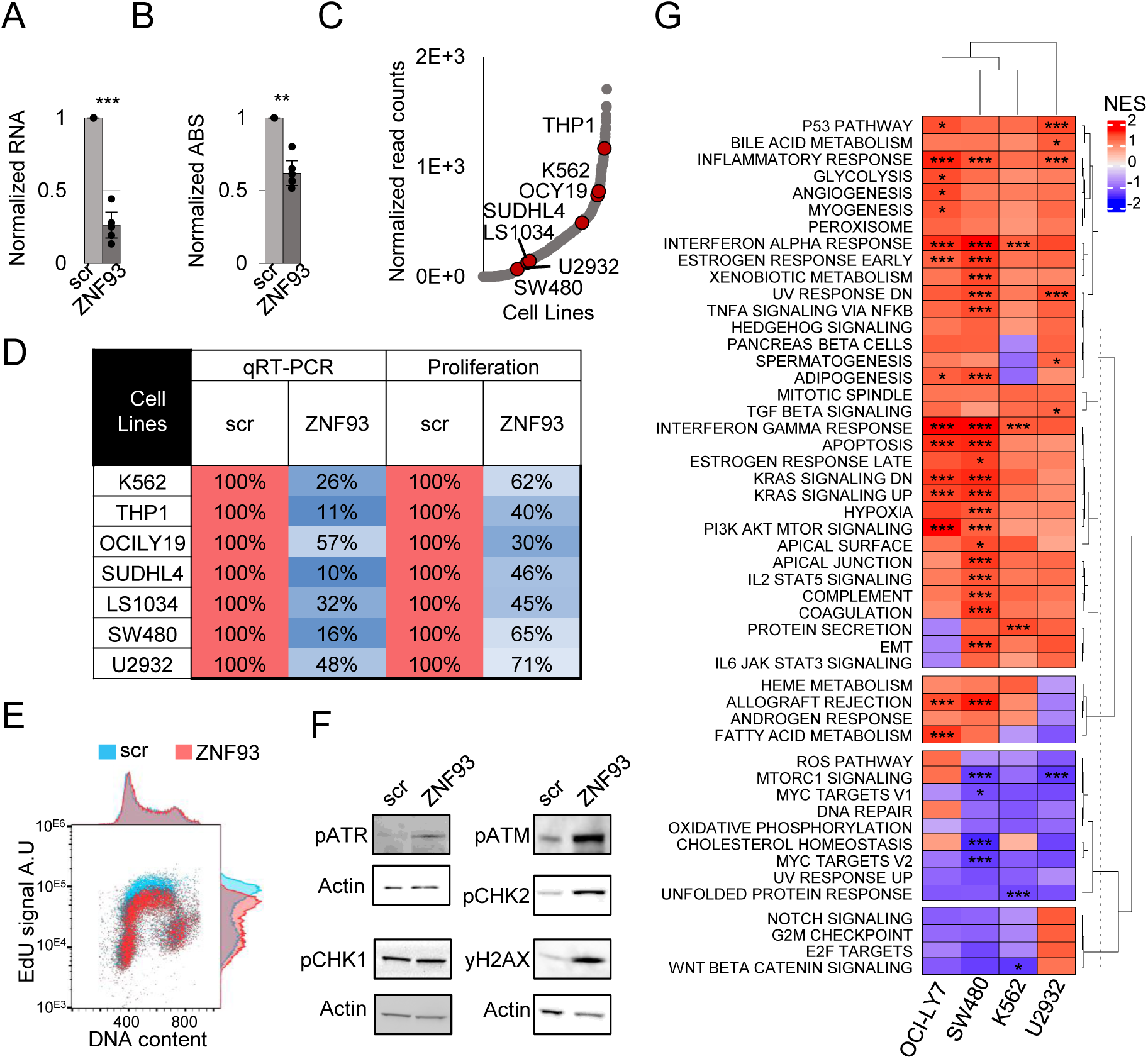
ZNF93 depletion reduces proliferation in cancer cells. A. *ZNF93* expression levels (RT-qPCR) upon ZNF93 knockdown (KD) in K562 cells or scrambled (scr) (n = 6, t.test). B. Metabolic activity as a surrogate for proliferation of K562 cells upon *ZNF93* KD (n = 6, t.test). C. *ZNF93* expression in cancer cell lines from (https://sites.broadinstitute.org/ccle/). Cell lines used in this study are highlighted in red. D. *ZNF93* KD efficiency and associated effects on cell proliferation across cell lines. E. Flow cytometry analysis of the replication signal (EdU incorporation intensity, y-axis) and DNA content (DAPI staining, x-axis) upon *ZNF93* KD in K562 cells. F. pATR, pATM, pCHEK2, pCHEK1, and γH2AX signals upon *ZNF93* KD in K562 cells. G. Heatmap plot of Normalized Enrichment Score (NES) of Hallmark signatures derived from RNA-seq data upon ZNF93 KD in the OCI-LY7, SW480, K562 and U2932. Asterisks indicate significant enrichment compared to control conditions.

These proliferation defects concurred with reduced incorporation of the thymidine analog EdU into newly synthesized DNA in S-phase (Fig. 3E, S3C), implying replication defects and activation of the replication and DNA damage checkpoints indicated by increased phosphorylation levels of ATR, CHK1, ATM, CHK2 and H2AX (Fig. 3F). Transcriptomic analysis of *ZNF93* KD cells by RNA-seq revealed that pathways elicited upon DNA damage and replication stress, including p53 signaling, apoptosis and inflammation were activated (Fig. 3G, S3D).

Together, these results show that ZNF93 depletion induces replication stress, activates the DNA damage response and inflammation, which result in stalling proliferation in multiple cancer cell lines.

### ZNF93 depletion modestly de-represses FLYL1, and fails to induce detectable changes in L1-encoded protein expression

We observed that ectopic overexpression of ORF2p, but not a catalytically dead endonuclease (dEN) mutant, recapitulated the phenotypes observed upon ZNF93 depletion. Confirming previous results (4), ORF2p overexpression led to reduced K562 proliferation (Fig. S4A, B), impaired EdU incorporation in S-phase (Fig. S4C) and pCHK1/2 accumulation (Fig. S4D). Given the similarity between the DNA damage response and replication stress phenotypes induced by *ZNF93* KD and ORF2p overexpression, we investigated whether the former resulted from an accumulation of ORF2p upon FLYL1^EN+^ de-repression. To this end, we characterized the epigenomic and transcriptional changes occurring at FLYL1^EN+^ loci upon *ZNF93* KD. CUT&Tag experiment revealed only modest reductions in H3K9me3 - the repressive heterochromatin mark instigated by the KZFP/TRIM28 axis - at ZNF93-bound FLYL1^EN+^ promoters, with many FLYL1^EN+^ loci remaining unaffected (Fig. 4A). RNA-seq analyses revealed that *ZNF93* KD induced FLYL1 upregulation to a degree that varied between different cell lines (Fig. 4B, S4E). However, this was not limited to ZNF93 L1 targets, as L1Hs and L1PA2 integrants also exhibited increased transcription (Fig. 4B, S4E). We sought to determine whether FLYL1 upregulation resulted in increased intracellular levels of the genotoxic, L1-encoded ORF2p. We successfully detected L1-encoded endogenous ORF1p by western blot, but found its expression to be unaffected by *ZNF93* KD (Fig. 4C). In spite of multiple attempts, we failed to detect endogenous ORF2p, whether in control or in *ZNF93* KD cells, consistent with the minute levels at which ORF2p is expressed relative to ORF1p (42, 43). Thus, we could not firmly establish that the observed DNA damage and replication stress observed upon ZNF93 depletion resulted from the uncontrolled expression of L1 ORF2p.

**Figure 4:**
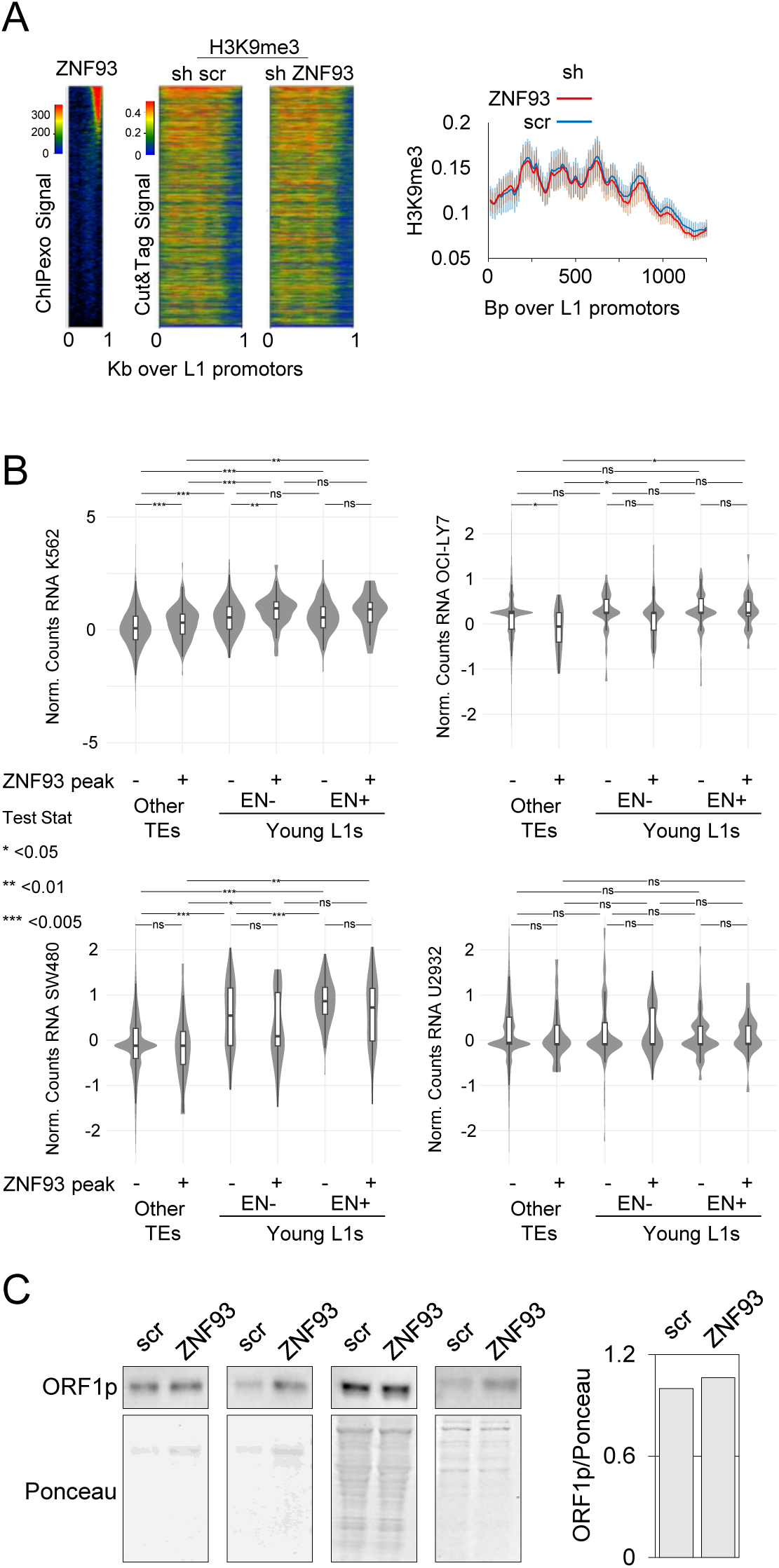
ZNF93 depletion modestly de-represses FLYL1, but fails to induce detectable changes in L1-encoded protein expression. A. Left: ZNF93-HA ChIPexo signal upon OE in HEK293T cells (Imbeault *et al.*, 2017) and H3K9me3 ChIP-seq signal upon *ZNF93* KD in K562 cells across FLYL1^EN+^ promoters. Right: H3K9me3 signal across the meta-FLYL1^EN+^ sequence upon *ZNF93* KD in K562 cells. B. Normalized RNA counts across transposable elements (TEs) upon *ZNF93* KD in K562, OCI-LY7, SW480, and U2932. Statistical test: two-tailed t-test. C. ORF1p signal upon *ZNF93* KD in K562 cells. Quantification relative to Ponceau S Staining on the right.

### ZNF93 is a critical regulator of *APOBEC3B* expression

Although ZNF93 primarily binds to L1 5’UTRs, it is also modestly enriched at some gene promoters, suggesting a potential role in regulating gene expression *via* promoter occupancy (Fig. S5A). We thus examined transcriptional changes occurring upon ZNF93 knockdown by combining RNA-seq data from *ZNF93* KD and *ZNF93* overexpression (OE) experiments in K562 cells (Fig. S5B), focusing on changes affecting genes with a ZNF93 ChIPseq peak within 50 kb window of their transcription start sites (TSS). This analysis identified the gene encoding APOBEC3B—a cytidine deaminase known to act as a DNA editor and inducer of replication stress and DNA damage—as a top ZNF93-regulated target. *APOBEC3B* was significantly upregulated upon *ZNF93* KD and downregulated upon *ZNF93* OE, and further displayed a prominent ZNF93 binding peak at its TSS (Fig. 5A).

**Figure 5:**
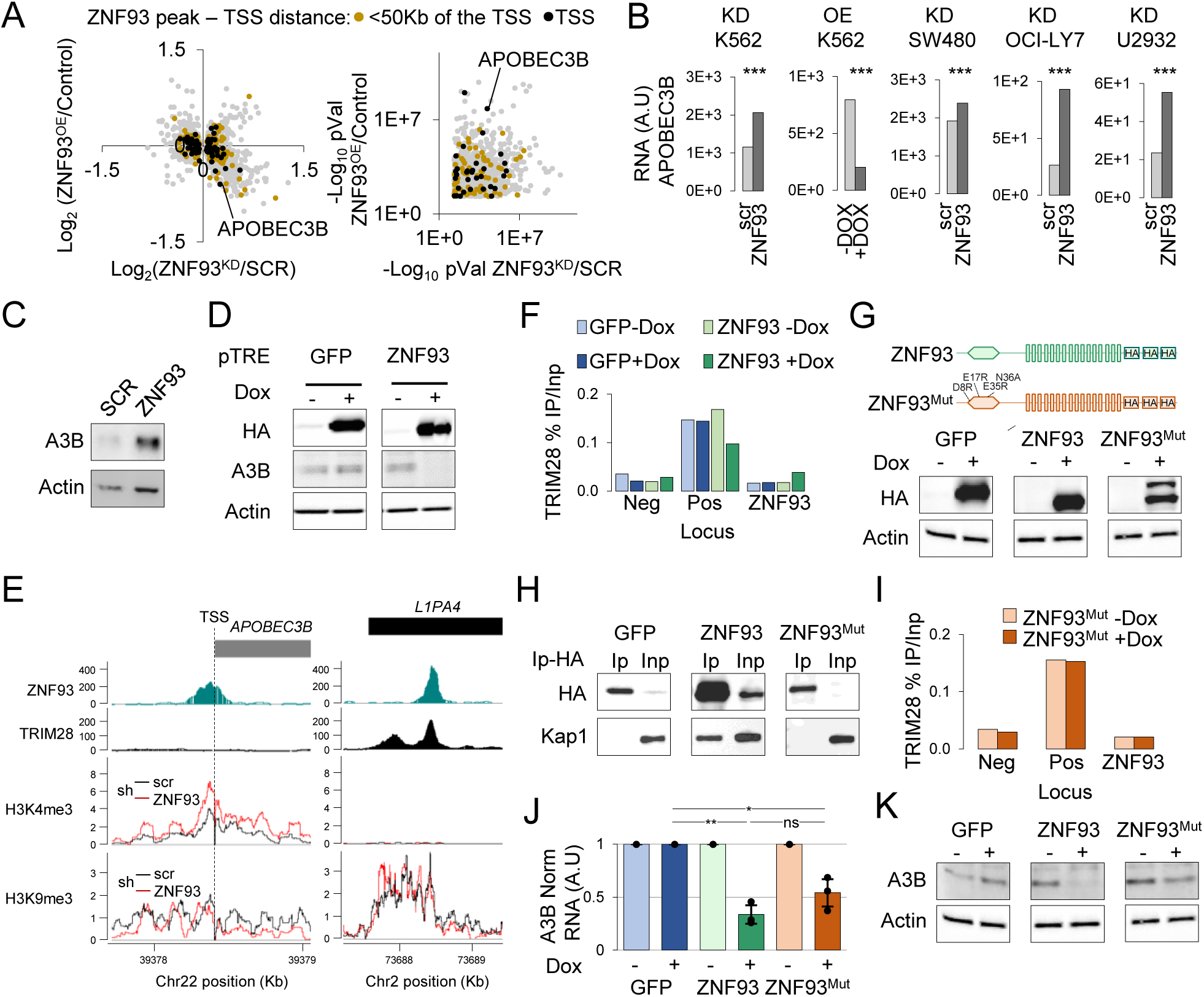
ZNF93 is a critical regulator of APOBEC3B expression. A. Gene expression changes upon *ZNF93* KD (x-axis) and overexpression (OE) (y-axis) in K562 cells. Orange dots indicate ZNF93 peaks within 50 Kb of the transcription start site (TSS), while black dots indicate ZNF93 peaks overlapping the TSS. Left: fold changes relative to control. Right: -Log10 p-value. B. *APOBEC3B* expression levels after *ZNF93* KD in K562, U2932, SW480, and OCI-LY7 cells, as well as five days after *ZNF93* OE induced by doxycyclin in K562 cells. C. APOBEC3B signal six days after *ZNF93* KD in K562 cells. Statistical test: Differential expression. D. APOBEC3B signal five days after OE of GFP-HA or ZNF93-HA in K562 cells. E. Integrated Genome Browser (IGB) screenshots showing the ZNF93 ChIP-exo track in 293T cells (Imbeault *et al.*, 2017), TRIM28 ChIP-seq in H1 cells (Castro Diaz *et al.*, 2014), and H3K9me3 and H3K4me3 CUT&TAG signals after *ZNF93* KD in K562 cells (sum of four independent replicats), at the *APOBEC3B* TSS and at one representative L1PA4. F. ChIP–qPCR analysis of TRIM28 binding at the *APOBEC3B* TSS after OE of GFP or ZNF93 in K562 cells. Means from two independent experiments are shown. Neg and Pos are negative and positive control regions for TRIM28 binding. G. Top: mutagenesis of ZNF93 (ZNF93^Mut^) with four amino acid substitutions in the KRAB domain, abrogating TRIM28 recruitment. Bottom: HA signal after OE of HA-tagged ZNF93, ZNF93^Mut^, or GFP in K562 cells. H. Co-immunoprecipitation of TRIM28 with HA-tagged proteins as described in Fig. 5G. HA was immunoprecipitated from K562 cell lysates six days after OE of ZNF93, ZNF93^Mut^, or control (GFP). I. ChIP–qPCR analysis of TRIM28 binding at the *APOBEC3B* TSS after OE of HA-tagged ZNF93, ZNF93^Mut^, or GFP in K562 cells. Means from two independent experiments are shown. J. RT-qPCR analysis of *APOBEC3B* expression five days after OE of HA-tagged ZNF93, ZNF93^Mut^, or GFP in K562 cells. Means and SD from three independent experiments are shown. *p < 0.05 (two-tailed t-test). K. APOBEC3B signal after OE of HA-tagged ZNF93, ZNF93^Mut^, or GFP in K562 cells.

Integrating RNA-seq data from additional cell lines (U2932, OCY7 and SW480) (Fig. S5C) revealed that *APOBEC3B* was in fact the only gene behaving as a strict ZNF93 target, being consistently upregulated across all *ZNF93* KD conditions (Fig. 5B, S5C). Accordingly, *ZNF93* KD led to a dramatic increase in APOBEC3B protein levels, which were conversely reduced in *ZNF93* OE K562 cells (Fig. 5C, D).

We next assessed whether the ZNF93-*APOBEC3B* regulatory dependency correlated with epigenetic changes at the *APOBEC3B* promoter. There was surprisingly no clear H3K9me3 peak at the *APOBEC3B* ZNF93 binding site (Fig. 5E), where TRIM28 was not recruited, in sharp contrast with what observed at FLYL1 promoters (Fig. 5E, 5F). However, there was an increase in the activation mark H3K4me3 at the *APOBEC3B* promoter in *ZNF93* KD K562 cells (Fig. 5E). This strongly suggested that the KZFP regulated the cytidine deaminase by non-conventional mechanisms.

To test this hypothesis, we mutated ZNF93 at residues of its KRAB domain previously determined to be essential for TRIM28 recruitment (Fig. 5G) (44). Co-immunoprecipitation confirmed that the resulting ZNF93^mut^ indeed failed to associate with the master corepressor (Fig. 5H, I). Nevertheless, upon overexpression this mutated protein still silenced *APOBEC3B* expression, as verified at both RNA and protein levels, albeit slightly less efficiently than its wild-type counterpart (Fig. 5J, K). Therefore, ZNF93 represses *APOBEC3B* expression largely through TRIM28-independent mechanisms.

### ZNF93 protects proliferating cells from APOBEC3B-induced replication stress

APOBEC3B expression is known to increase in response to cancer cell proliferation and genotoxic stress, and in return to amplify replication stress (26–33). Having established that ZNF93 silencing upregulates APOBEC3B, we hypothesized that ZNF93 may limit replication stress in proliferating cells by repressing the cytidine deaminase. To test this hypothesis, we characterized the impact of ZNF93 on cellular responses to the genotoxic stressor hydroxyurea (HU). As previously observed (31), HU treatment led to APOBEC3B induction and increased γH2AX levels, indicative of DNA damage and replication stress. This response was markedly reduced by *ZNF93* overexpression (Fig. 6A). Furthermore, while *ZNF93* OE could not alleviate the HU-induced replication block (Fig. S6A), it allowed for a swifter return to proliferation following HU washout, similar to what observed upon *APOBEC3B* knockdown (Fig. 6B), whereas the opposite was observed upon *APOBEC3B* overexpression (Fig. 6B).

**Figure 6:**
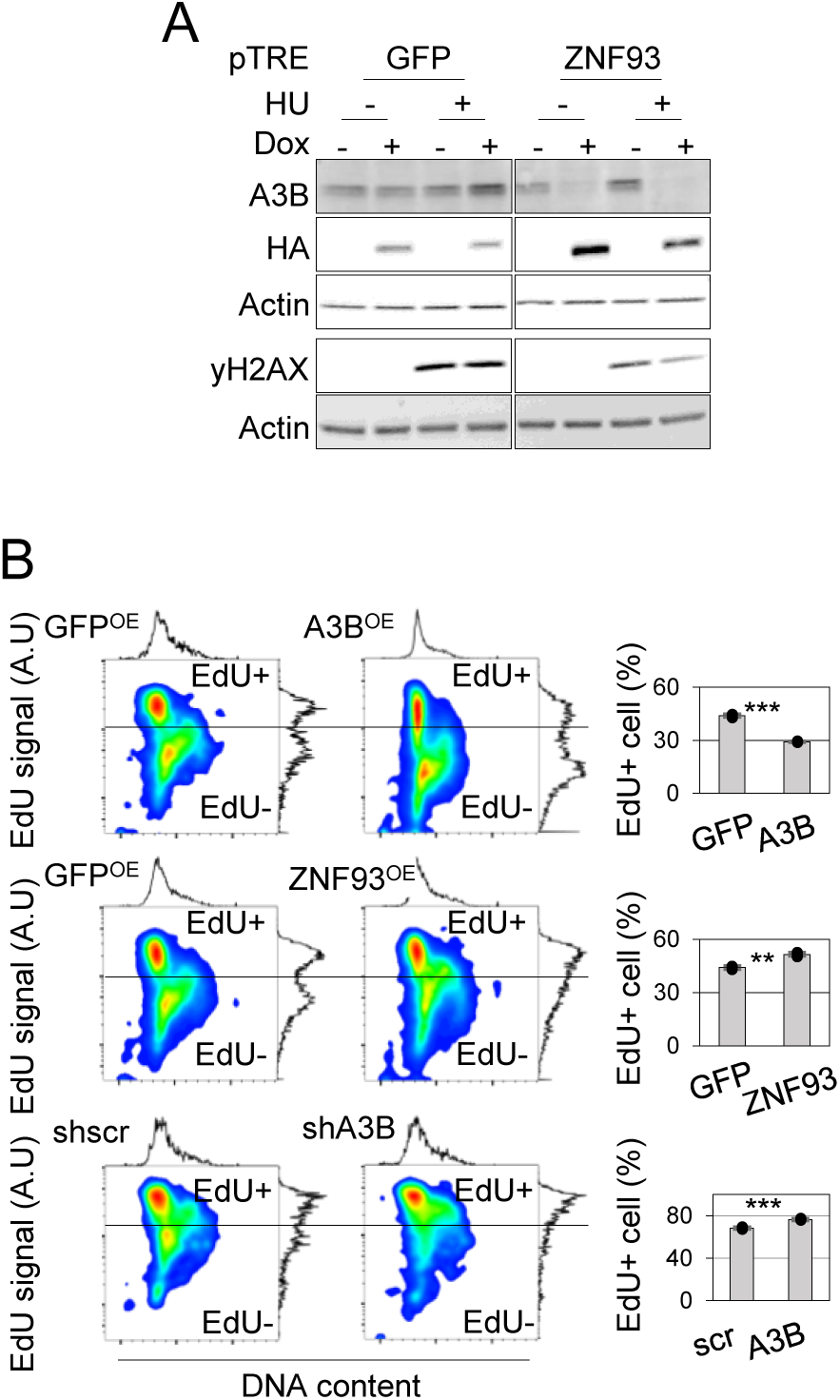
ZNF93 protects proliferating cells from APOBEC3B-induced replication stress. A. APOBEC3B and γH2AX signal three days after induction of GFP-HA or ZNF93-HA, followed by hydroxyurea (HU) treatment for 20 hours in K562 cells. B. Left: EdU-FACS analysis three days after induction of GFP-HA, APOBEC3B-HA, or ZNF93-HA, or after shRNA-mediated depletion of APOBEC3B, followed by HU treatment for 20 hours and an eight-hour recovery period in K562 cells. Right: Quantification of EdU+ cells. Statistical significance: **p < 0.01; ***p < 0.001 (two-tailed t-test).

## DISCUSSION

Our study identifies the primate-specific KZFP ZNF93 as a pivotal guardian of genome stability. While ZNF93 has been recognized for its role in silencing TEs, particularly L1 retrotransposons (6), we provide evidence that it more broadly mitigates replication stress in proliferative or genotoxic contexts by also repressing the expression of the APOBEC3B cytidine deaminase.

APOBEC3B, a primate-restricted member of the APOBEC family of viral restriction enzymes, uniquely locates to the nucleus, where it restricts both retrotransposons and viruses through DNA editing. In addition, APOBEC3B has emerged as a critical regulator of secondary structures known as R-loops during DNA replication (45). Fittingly, APOBEC3B exhibits a robust cell cycle-rhythmic expression pattern, reaching maximal expression during the S phase (40). Thus, APOBEC3B, similar to SAMHD1 and other innate immune related proteins, stands at the crossroads of immunity and DNA replication (46–48), bridging the oft-mentioned divide separating fast-evolving, environment-interacting genes – e.g. immune genes – from ultra-conserved genes involved in ubiquitous processes such as the cell cycle (49). Related to this, we have recently reported that surprisingly many KZFPs – given their recent evolutionary emergence – regulate cell cycle-oscillating genes via promoter binding, suggesting unappreciated levels of lineage specificity in the regulation of ubiquitous cellular processes (40). The existence of a wholly primate-specific ZNF93-APOBEC3B regulatory axis further indicates that fundamental biological processes are not solely carried out by deeply conserved genes.

APOBEC3B, together with APOBEC3A, has been identified as a potent host mutagen in multiple cancer cell types, contributing to tumor heterogeneity, evolution and resistance to therapy (31, 37, 38, 50, 51). Yet, unabated levels of APOBEC3B-mediated mutagenesis may result in unsustainable, fatal levels of replication stress and DNA damage (52). The high correlation observed between ZNF93 expression and cell proliferation across most cancer types suggests that by repressing APOBEC3B, ZNF93 partakes in this trade-off. The ZNF93-APOBEC3B regulatory dependency may therefore have important implications for tumor progression. Supporting this, the only cancer type where ZNF93 expression anti-correlates with proliferation according to TCGA (BH-adj. p < 0.05, rho < 0) is cervical squamous cell carcinoma (CESC). CESC is frequently caused by human papillomavirus (HPV) infection, which APOBEC3B participates in restricting. This antiviral action results in the collateral accumulation of mutations, thereby favouring cancer initiation. Accordingly, we noted that compared to HPV-negative tumours, HPV-positive tumours, which express high levels of *APOBEC3B* (53), simultaneously exhibit low *ZNF93* expression (Fig. 7). It is thus tempting to speculate that in cancer cells, regulating ZNF93 expression may be adaptative and partake in balancing APOBEC3B-mediated mutagenesis against APOBEC3B-induced replication stress. Interestingly, ZNF93 has been found to confer human chondrosarcoma cells with resistance to the inducer of replication stress and DNA damage ET-743 (Trabectedin; Yondelis®), (54) an effect potentially linked to ZNF93-mediated repression of APOBEC3B.

**Figure 7:**
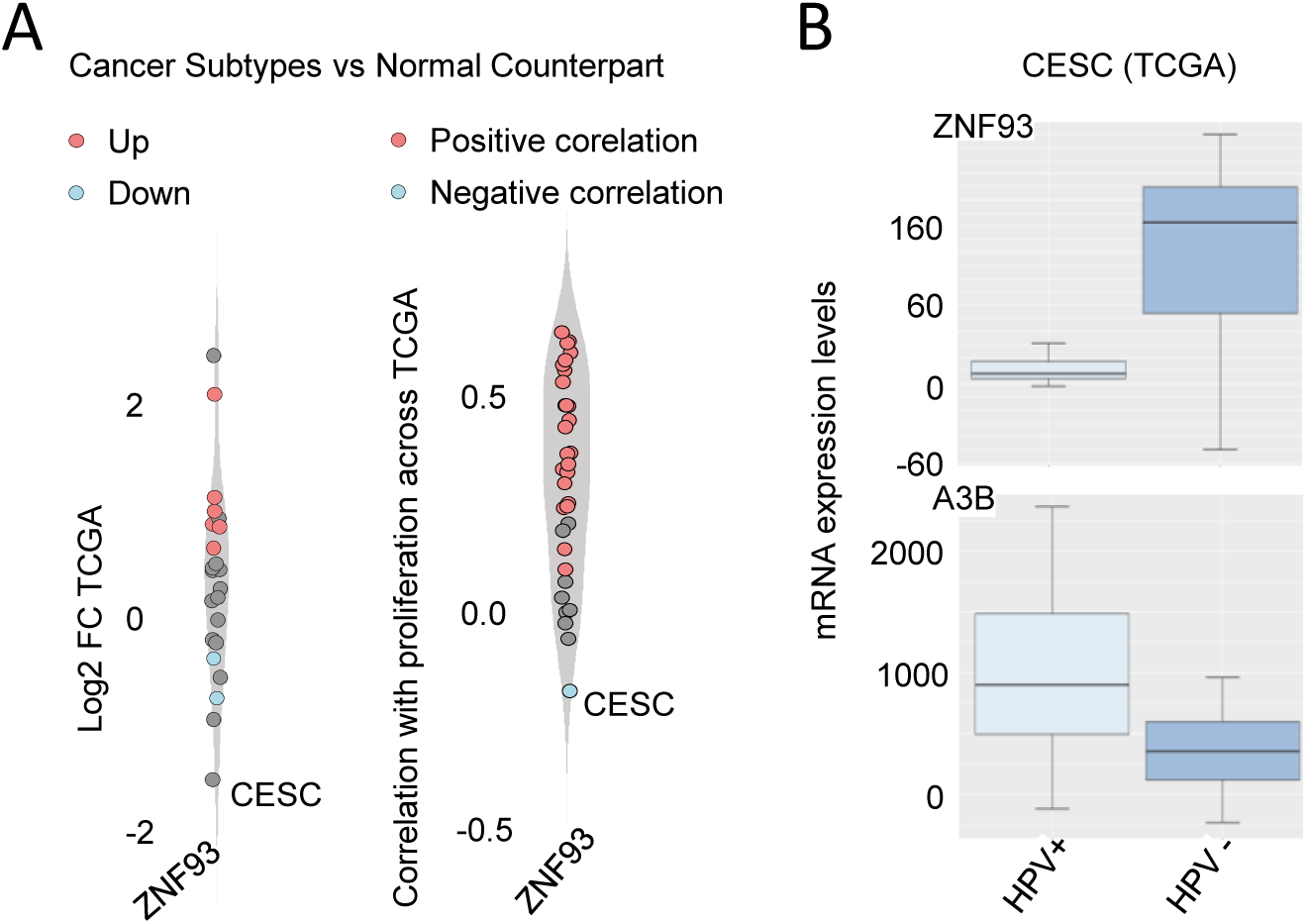
ZNF93 is downregulated in HPV-positive cervical cancer samples. A. Left: differential *KZFP* expression between cancer and normal samples (23 cancer subtypes selected). Median Spearman correlation coefficients and log2FC were compared across subtypes using Wilcoxon’s two-tailed test. Right: Spearman rank correlation (t-test, adj. p < 0.05) between *KZFP* expression and a 167-gene proliferation signature across 33 TCGA cancer subtypes. Red: significant correlation; Blue: significant anticorrelation. CESC position is annotated B. From Salnikov *et al.*, 2022. Comparison of *ZNF93* and *APOBEC3B* mRNA expression across HPV-positive (HPV+), HPV-negative (HPV-), in the TCGA cervical (CESC) cohorts. Expression data (Level 3) was obtained from the Broad Genome Data Analysis Center’s Firehose server. HPV status annotation follows Gameiro *et al*., 2017.

ZNF93-mediated regulation of APOBEC3B appears partially independent of TRIM28, despite robust TRIM28 recruitment at ZNF93-bound TEs. Similar dissociations between TRIM28 and KZFP-mediated promoter regulation have been observed previously (55), suggesting that KZFPs may employ distinct mechanisms at gene promoters compared to TEs. Future work should explore whether steric hindrance, for example competition with the binding of specific transcription factors, underlies this TRIM28-independent regulatory activity, and the mechanisms responsible for the failure of canonical KZFPs to recruit the co-repressor at some but not other genomic loci.

Questions remain regarding the impact of ZNF93-mediated control of FLYL1. Upon observing the genotoxic effect of ZNF93 depletion, we first thought that it resulted from the production of ORF2p endonuclease by ZNF93-repressed FLYL1 integrants. We speculated that the preservation, in some cases for more than 20 million years, of the toxic endonuclease-coding potential of thousands of integrants dispersed throughout the genome allowed the cell to use them as sentinels watching for loss of epigenetic control. However, we only observed a mild upregulation of FLYl1s in the ZNF93 knockdown cancer cell lines tested here, with minimal changes in H3K9me3 on their 5’UTR, no discernable elevation in ORF1p levels and no detectable ORF2p. While this could be attributed to the effect in these cells of other KZFPs binding to the promoter of these L1 integrants, such as ZNF649 and ZNF765, we cannot formally exclude that minute amounts of highly active endonuclease, which is physiologically produced at levels between 200 and 500 lower than ORF1p, were partly responsible for part of the observed phenotype, nor that this protein plays a more prominent surveillance role in other settings.

Irrespectively, our data support a model whereby KZFPs and their genomic targets act as rheostats of genomic instability in cancer cells, with their combined actions likely exerting broad influences on DNA damage, replicative stress, DNA editing and other forms of mutagenesis, inflammation, immune escape and resistance to some therapies. As recently demonstrated for ZNF417 and ZNF587, the upregulation of which protects diffuse large B cell lymphoma cells by minimizing HERVK-induced genotoxicity, the ZNF93-APOBEC3B regulatory axis represents a primate-restricted determinant of replication stress, underscoring an underappreciated level of lineage-specificity in the management of genome integrity.

## MATERIAL AND METHODS

### L1s characterization

Elements are annotated as full length if the calculated length using hg19 annotation is > 5900 bp. Over full length L1s, ORF2p were detected using ORF detection (56) with the following parameters : Minimum nucleotide size of ORF to report : 30, Maximum nucleotide size of ORF to report : 10000, What to output : Translation of regions between START and STOP codons, All START codons to code for Methionine : true, Circular sequence : false, Find ORFs in the reverse complement, true, Number of flanking nucleotides to output : 0, Output sequence file format : FASTA (m).

Amongst the detected ORFs, L1-ORF2p were detected using the following script in R:

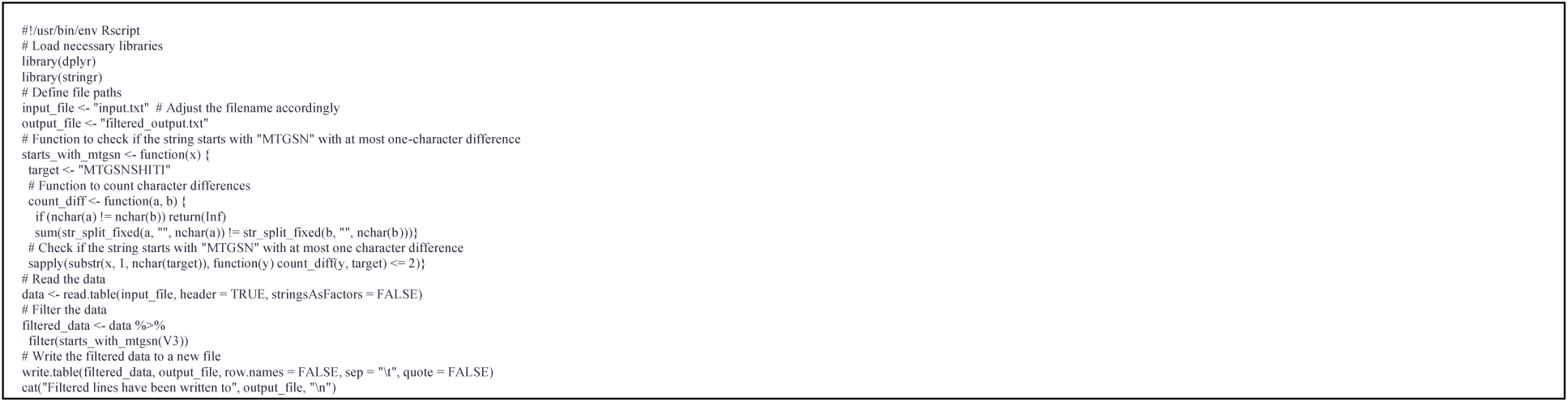

L1s are annotated as FLYL1^EN+^ if > if > 230 aa and FLYL1^EN-^ if not. Output: **Table S1**.

### Enrichment of KZFPs on repeats

Enrichment for KZFPs over L1s were extracted from (7). Enrichments were computed with pyTEnrich available at URL https://github.com/alexdray86/pyTEnrich. Association with pVal < 0.05 are considered significant and used for further analysis.

### Average profiles of KZFPs on L1s regions

Matrixes were generated using deeptools 2 (57): computeMatrix with the L1s coordinates (**Table S1**) (-R) and various Bigwigs (-S)”; and the following options -bs 50 –sortRegions keep –sortUsing mean –averageTypeBins mean –outFileSortedRegions”. Then, matrices were used to perform average profiles using plotProfile software from Deeptools2 using following options “–averageType mean”. The following script was used :

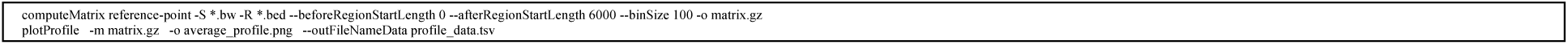

### TCGA Analysis

Analysis as in (40) with the following tables **Table S2**, **Table S3**. Represented with the following script:

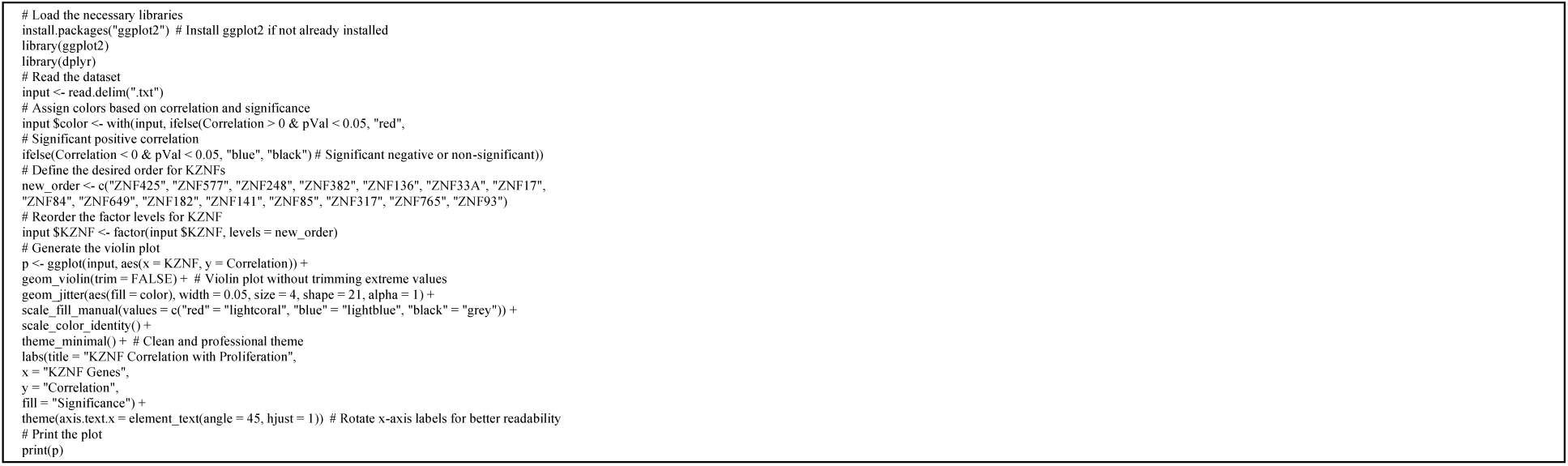

### CD4+T cell dataset

The datasets to analyse RNAseq and KAP1 ChIPseq are public (41). Gene Ontology enrichment was performed with the following script:

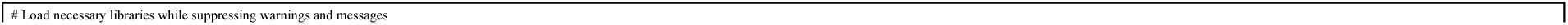
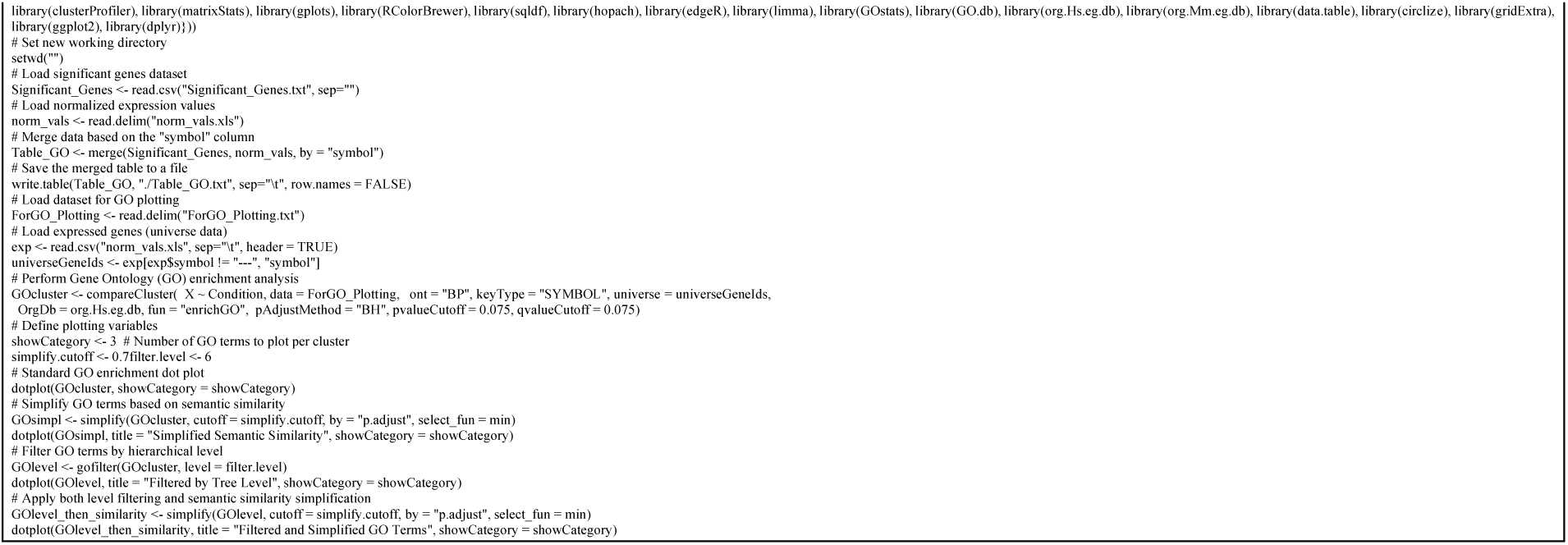

The list of genes associated with proliferation : as in (40).

### Cell culture

OCI-Ly7, OCI-Ly19 and U2932 human lymphoma cell lines were obtained from DSMZ – German Collection of Microorganisms and Cell Cultures, Braunschweig, Germany (www.dsmz.de). SUDHL4 were kindly supplied by Elisa Oricchio (EPFL, Swiss Institute for Experimental Cancer Research) and typed using short tandem repeat (STR) profiling by Microsynth cell line authentication service. THP1 were kindly supplied by Caroline Arber (Faculty of Biology and Medicine, University of Lausanne (UNIL). Immuno-Oncology Service (ION)). K562, LS1034, and SW480 cell lines were obtained from the American Type Culture Collection (ATCC). SW480 were cultured in L15 medium (Sigma) supplemented with 10% fetal bovine serum and 1% penicillin-streptomycin. K562, THP1, OCI-LY19, U2932, SUDHL4, and LS1034 cells were grown in RPMI 1640 (Gibco) with 10% fetal bovine serum and 1% penicillin-streptomycin. OCI-Ly7 cells were grown in IMDM (Gibco) with 20% fetal bovine serum (FBS) and 1% penicillin-streptomycin. All cell lines were grown at 37°C and 5% CO2.

### Lentivector production and stable gene expression

Cells overexpressing HA-tagged constructs were generated as described in (58). In short, cDNAs for HA-tagged constructs were codon-optimized, synthesized using the GeneArt service from ThermoFisher, and cloned into the doxycycline-inducible expression vector pTRE-3HA. Stable cell lines were generated using lentivector transduction of mycoplasma free HEK293T cells as described on http://tronolab.epfl.ch.

### Knockdown (KD) Protocol

Cells were transduced with lentivector particles expressing shRNA targeting the gene of interest (shZNF93: TRCN0000018246, Sequence: 5’-GCCTTAGTAGACATGAGATAA-3’) or scrambled sequence (shSCR – Sequence: 5’-CAACAAGATGAAGAGCACCAAG-3’). Cells were seeded at a density of 1x10⁵ cells/mL and transduced. After 48 hours, cells were treated with 1 µg/mL puromycin for 3 days to select for successfully transduced cells. After selection, cells were replated at 1x10⁵ cells/mL and collected for further analyses at various time points up to 10 days post-transduction.

### RT-qPCR

Total RNA extraction was performed using the NucleoSpin RNA plus kit (Machery-Nagel) according to the manufacturer’s recommendations. cDNA synthesis for qPCR was conducted using the Maxima H minus cDNA synthesis master mix (ThermoScientific). Real-time quantitative PCR was performed using PowerUp SYBR Green Master Mix (ThermoScientific) and run on a QuantStudioTM 6 Flex Real-Time PCR System. The following primers were used : TBP, ALAS1, ZNF93, APOBEC3B

### Cell proliferation assays for KD experiments

Cell proliferation was assessed using the PrestoBlue™ and crystal violet assays. K562, U2932, OCI-LY19, THP1, and SUDHL4 cells were plated five days post-LV transduction and after three days of puromycin selection. Cells (100 µL) were seeded in duplicate 96-well plates at 20,000 cells/well. PrestoBlue™ (10 µL) was added, and after a 3-hour incubation at 37°C, absorbance was measured at 570 nm, with background subtraction at 600 nm. Absorbance was measured again after four days to assess proliferation. For LS1034 and SW480 cells, plating occurred under the same conditions at 20% confluence in plates and flasks, respectively. For SW480 and LS1034, proliferation was evaluated using crystal violet staining. Five days post-LV transduction Cells were seeded in flasks and multi-well plates respectively at 20%, allowed to growth for 3 days, and washed twice with PBS after treatment. Fixation was performed with 100% methanol for 10 minutes at room temperature, followed by air drying. Cells were then stained with 0.5% (w/v) crystal violet in 20% methanol for 10–15 minutes with gentle rocking. Excess stain was removed by washing three to four times with distilled water. Plates were air-dried, and for quantification, the stain was solubilized with 10% acetic acid, incubated for 10–15 minutes, and absorbance was measured at 590.

### EdU DNA synthesis monitoring flow cytometry

K562 cells were pulse-labeled with 10 µM 5-ethynyl-2′-deoxyuridine (EdU) for 20 min and SW480 and LS1034 for 40 min and subsequently fixed with 2% formaldehyde for 30 min at room temperature (RT). EdU incorporation was detected using Click chemistry according to manufacturer’s instructions (Click-iT EdU Flow Cytometry Cell Proliferation Assay, Invitrogen). Cells were resuspended in 1x Phosphate Buffer Solution (PBS, BioConcept) with 1% bovine serum albumin (BSA), 2 µg/ml DAPI and 0.5 mg/ml RNase A for 30 min at RT and subsequently analyzed on a BD LSR II (Becton Dickinson, USA) flow cytometer, using BD FACSDivaTM software, and quantified using FlowJo single-cell analysis software (FlowJo, LLC).

### Western Blot

Proteins were extracted from cells using a homemade RIPA buffer (50 mM Tris-HCl, 150 mM NaCl, 1% NP-40, 0.1% SDS, 0.5% sodium deoxycholate, pH 7.4) and quantified using the BCA Protein Assay Kit (Pierce). Equal amounts of protein were separated on a 4-20% gradient pre-cast Tris-glycine gel from Thermo Fisher and transferred onto a nitrocellulose membrane using the iBlot 3 system from Thermo Fisher. The membrane was blocked with 5% non-fat milk in PBS-T (PBS containing 0.1% Tween-20) for 1 hour at room temperature, followed by overnight incubation at 4°C with primary antibody as followed : Anti-HA-Peroxidase (High Affinity, 50 mU/ml, Roche, ref: 12013819001), HRP-conjugated anti-actin antibody (1:5000 dilution, ThermoFisher, ref: MA5-15739-HRP), Phospho-Chk1 (Ser345) (133D3) Rabbit mAb 2348, Phospho-Chk2 (Thr68) (C13C1) Rabbit mAb 2197, Phospho-ATM (Ser1981) (D6H9) Rabbit mAb #5883, Phospho-ATR (Ser428) Antibody #2853 and Phospho-Histone H2A.X (Ser139) (20E3) Rabbit mAb #9718, APOBEC3B (E9A2G) Rabbit mAb #41494, ORF1p (D3W9O) Rabbit mAb #88701. in 5% non-fat milk in PBS-T. After three washes with PBS-T. membranes were incubated with goat anti rabbit antibody (1:5000Anti-rabbit IgG, HRP-linked Antibody #7074). After 3 wash, protein bands were visualized using ECL detection reagent (Advansta Inc, ref: K-12049-D50) and imaged using a chemiluminescent imaging system (Fusion FX from Vilber).

### Design of L1-ORF2p

cDNAs for HA-tagged constructs were codon-optimized, synthesized using the GeneArt service from ThermoFisher, and cloned into the doxycycline-inducible expression vector pTRE-3HA with the following sequences:

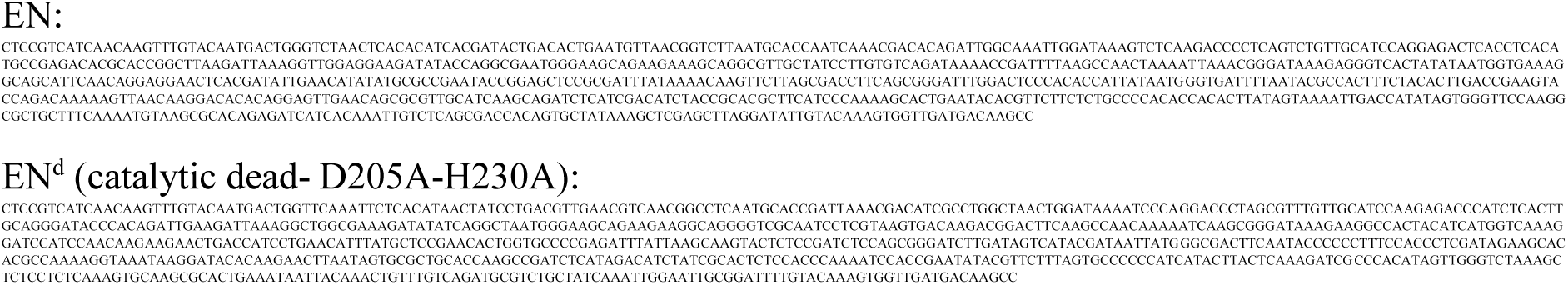

### Overexpression (OE) Protocol for EN and EN^d^ in K562 Cells

Doxycycline (Dox)-inducible overexpression was performed using a lentiviral system. K562 cells were transduced with lentiviral particles containing EN-3XHA constructs under the control of a tetracycline-inducible promoter. Cells were seeded at 1x10⁵ cells/mL and transduced. After 48 hours, cells were selected with 1 µg/mL puromycin for 3 days. Overexpression was induced by adding 1 µg/mL doxycycline to the culture medium for 3 days. Induction efficiency was validated by Western blot.

### Cell proliferation assays for OE experiments

The PrestoBlue™ assay was used to assess the proliferation of K562 cells. Following 5 days of lentiviral (LV) transduction and 3 days of puromycin selection, cells were either induced with doxycycline to express the transgene (+Dox) or left uninduced (-Dox). At each designated time point, 100 μL of cell culture was incubated with 10 μL of PrestoBlue™ reagent for 3 hours at 37°C. Absorbance was then measured immediately on a microplate spectrophotometer at 570 nm, with background absorbance at 600 nm subtracted for accuracy. Data normalization was conducted against the -Dox condition to account for baseline cell count variations.

### Cleavage Under Targets and Tagmentation (CUT&Tag)

CUT&Tag was performed as described (59) without modifications. For each mark, 150k cells were used per sample using the anti-H3K9me3 primary antibody (Active Motif, AB_2532132), the anti-H3K4me3 primary antibody (C42D8) and anti-rabbit IgG (Abcam, ab46540) secondary antibody. A homemade purified pA-Tn5 protein (3XFlag-pA-Tn5-Fl, addgene #124601) was produced and coupled with MEDS Oligos by the Protein Production and Purification of EPFL, as previously described. Purified recombinant protein was used at a final concentration of 700 ng/uL (1:250 dilution from homemade stock). Libraries were sequenced with 75 bp paired end on the NextSeq 500 (Illumina). Reads were aligned to the hg19 reference genome using bowtie2 (60). Only proper read pairs with MAPQ>10 were kept. Bigwig coverage tracks with the mean of replicate samples were generated using bedtools 2.27.168 (61) and deeptools 3.3.169 (57), and heatmap representations of the coverage signal were performed using computeMatrix function and plotHeatmap from deeptools 3.3.1. The following code was used:

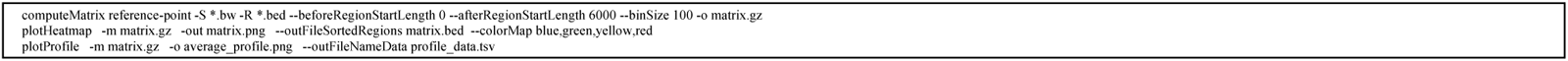

### RNA extraction

Total RNA extraction was performed using the NucleoSpin RNA plus kit (Machery-Nagel) according to the manufacturer’s recommendations.

### RNA-seq librairies and downstream analyses

RNA-seq libraries were prepared using the Illumina Truseq Stranded mRNA kit. Libraries were sequenced in 75 or 100 bp paired-end formats on the Illumina Hiseq 4000 and NovaSeq 6000 sequencers, respectively. RNA-seq reads were mapped to the hg19 human genome releases using hisat v2.1.060. Only uniquely mapped reads were used for counting over genes and repetitive sequence integrants (MAPQ > 10). Counts for genes and TEs were generated using featureCounts v2 and normalized for sequencing depth using the TMM method implemented in the limma package of Bioconductor. Counts on genes were used as library size to correct both gene and TE expression. For repetitive DNA elements, an in-house curated version of the Repbase database was used. Differential gene expression analysis was performed using Voom61 as implemented in the Limma package of Bioconductor62. P-values were adjusted for multiple testing using the Benjamini-Hochberg method. TEs were compared with the following scripts applied on **Table S3**:

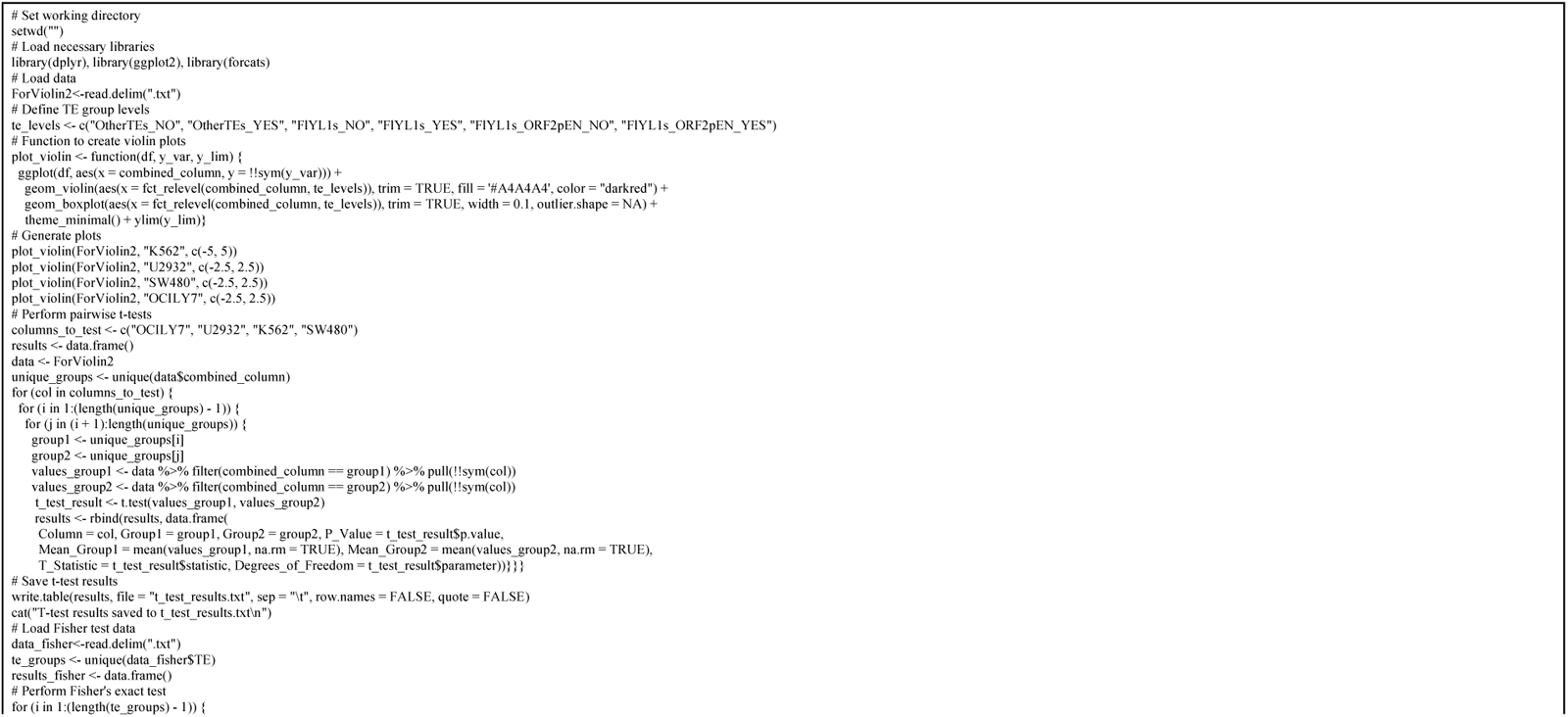
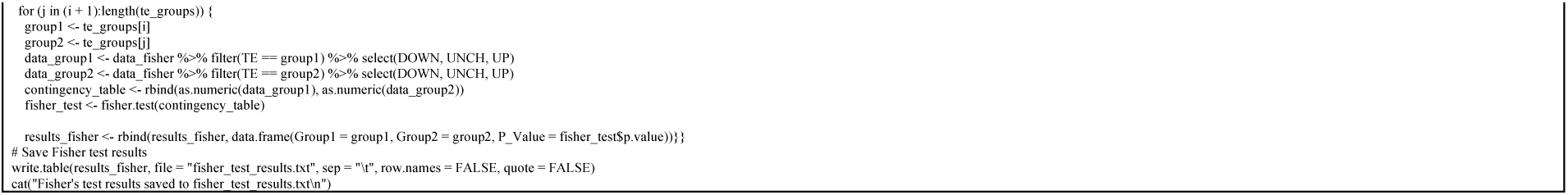

Gene Set Enrichment Analysis for ZNF93 KD was done using GSEA web interface (62, 63). The following option were used:

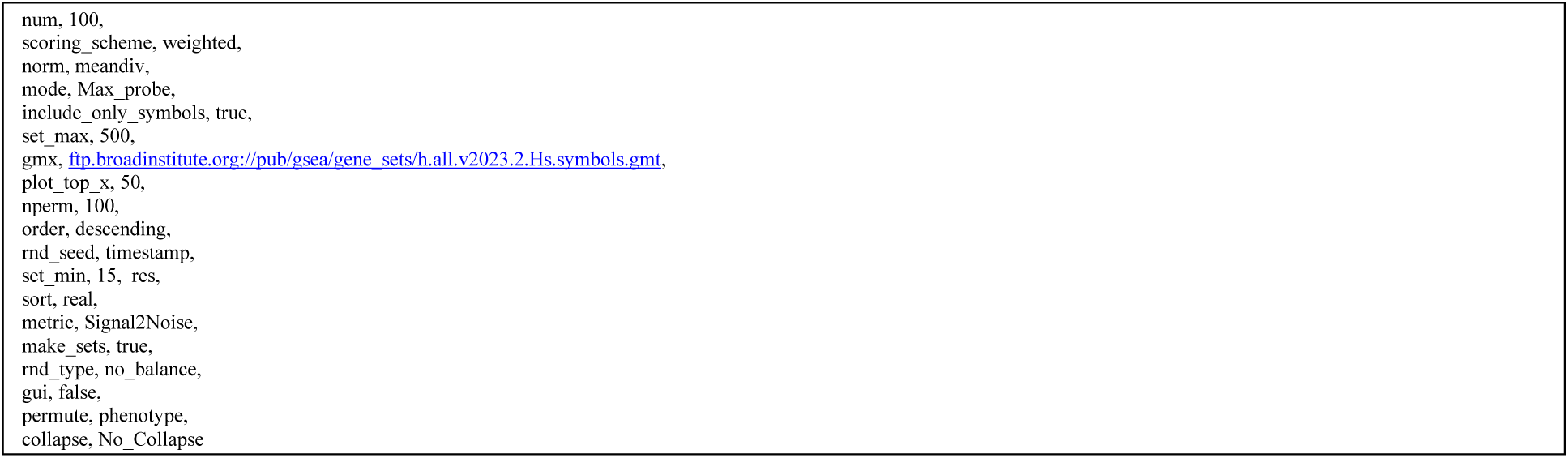

and the following script was used to build the heatmap:

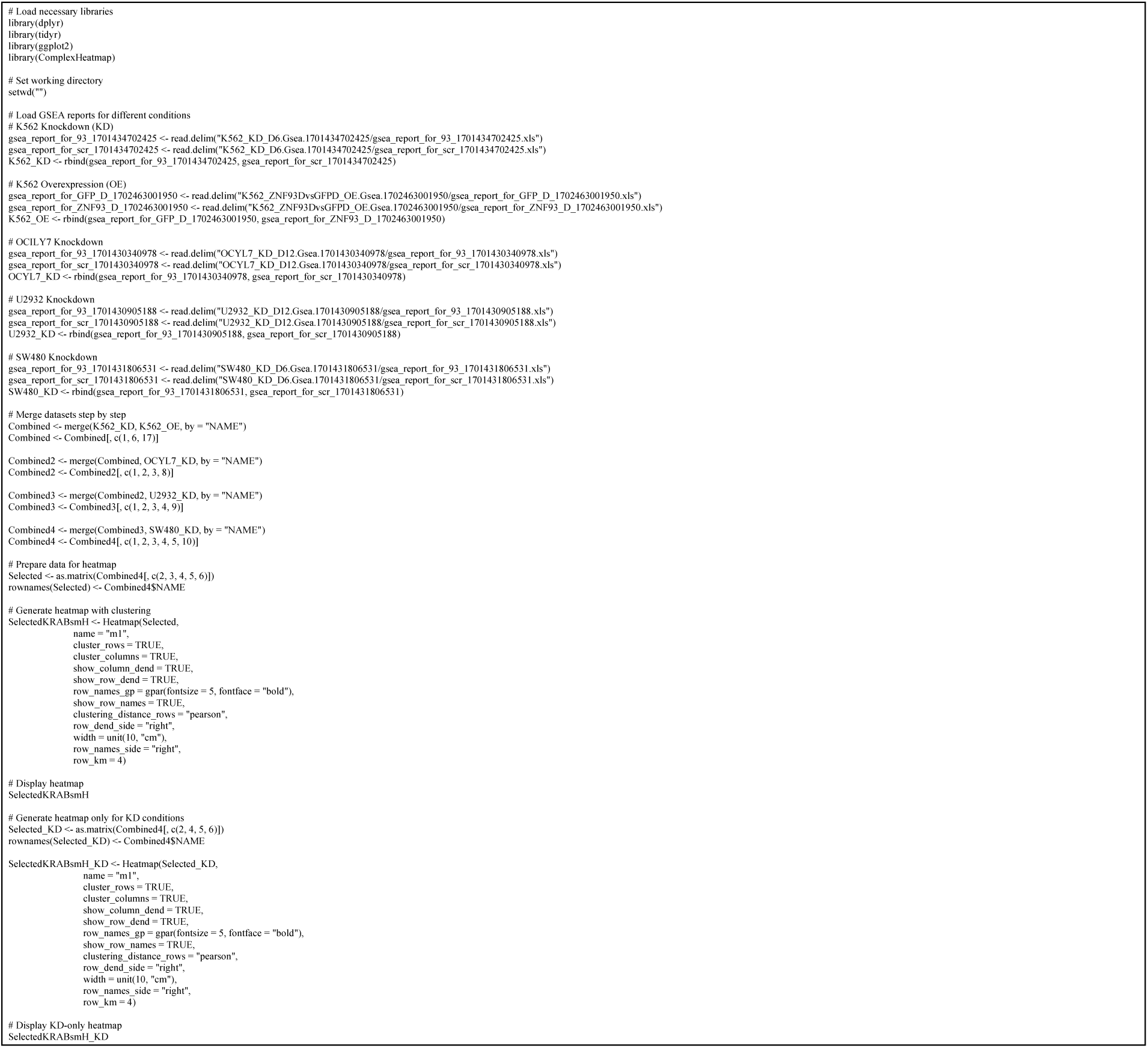

Genes were analysed for ZNF93-HA binding proximity (9) and intersection between datasets was computed using the following website : https://bioinformatics.psb.ugent.be/webtools/Venn/ and plotted with the following script:

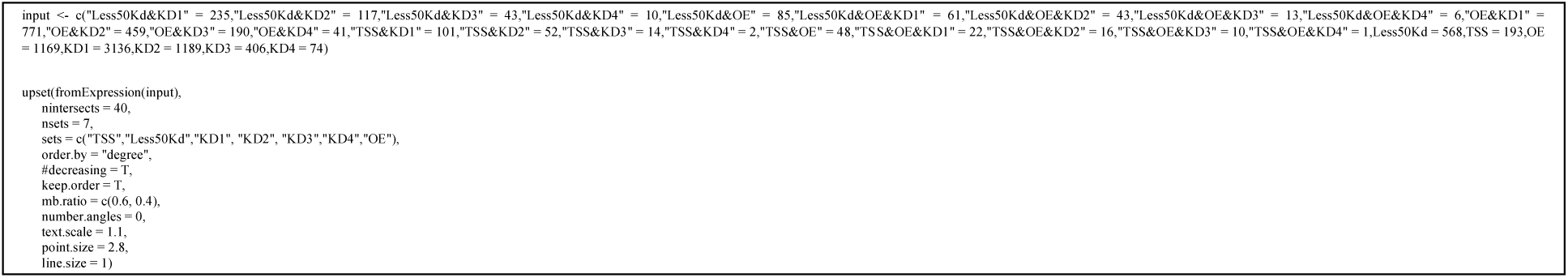

Gene analysis from the RNAseq including the result of the intersection between the datasets are reported Table S4.

### Overexpression (OE) Protocol for ZNF93 in K562 Cells

Doxycycline (Dox)-inducible overexpression was performed using a lentiviral system. K562 cells were transduced with lentiviral particles containing GFP-3XHA, LACZ-3XHA or codon optimized ZNF93-3XHA constructs under the control of a tetracycline-inducible promoter. Cells were seeded at 1x10⁵ cells/mL and transduced. After 48 hours, cells were selected with 1 µg/mL puromycin for 3 days. Overexpression was induced by adding 1 µg/mL doxycycline to the culture medium for 5 days. Induction efficiency was validated by Western blot.

### Generation of ZNF93^MUT^

ZNF93MUT-HA was generated by introducing four mutations into pTRE-ZNF93-HA via site-directed mutagenesis. The plasmid was digested with XhoI and NheI. Mutations were introduced using In-Fusion cloning with specific primers: Forward Primer [Insert sequence] and Reverse Primer [Insert sequence]. PCR was performed with high-fidelity polymerase, followed by gel purification and recombination with the linearized plasmid using the In-Fusion HD Cloning Kit (Takara Bio©) according to the manufacturer’s protocol. The product was transformed into DH5α E. coli cells, and colonies were screened by PCR. Positive clones were verified by Sanger sequencing, and confirmed ZNF93MUT-HA plasmids were propagated and purified for further use.

### Co-Immunoprecipitation (Co-IP)

K562 cells overexpressing HA-tagged ZNF93, ZNF93^MUT^, or GFP were harvested 72 hours post-doxycycline induction, washed with ice-cold PBS, and lysed in 500 µL of IP-WCL buffer (9.5 mL IP buffer base, 20 µL of 1M benzamidine, 100 µL of 0.1M PMSF, 500 µL of Igepal 10% CA-630, and protease inhibitors). Lysates were sonicated (2 x 5 seconds, Amplitude 0.30) and clarified by centrifugation at 5,000 rpm for 5 minutes at 4°C. The supernatant was collected for immunoprecipitation. HA-tagged proteins were immunoprecipitated by adding 25 µL of anti-HA magnetic beads (Pierce) per sample, which were resuspended in 800 µL of dynabuffer and washed once with 1 mL of dynabuffer and twice with 1 mL of IP-WCL buffer. Beads were incubated with 500 µL of lysate overnight at 4°C with gentle rotation. After incubation, beads were washed three times with 800 µL of wash buffer (9.5 mL IP buffer base, 20 µL of 1M benzamidine, 100 µL of 0.1M PMSF, 500 µL of Igepal 10% CA-630, and protease inhibitors). For elution, beads were incubated with 62.5 µL of 1X loading buffer, heated at 95°C for 10 minutes, and analyzed by SDS-PAGE. Proteins were transferred to membranes, blocked with 5% milk or BSA in PBS-T, and probed with anti-KAP1 antibody (ab10483) overnight at 4°C. After incubation with HRP-conjugated secondary antibody, signals were detected using chemiluminescence (Pierce). Input and were collected, treated with 12.5 µL of 5X loading buffer, heated, and analyzed in parallel to confirm the presence of KAP1 and HA-tagged proteins.

### Chromatin Immunoprecipitation followed by qPCR (ChIP-qPCR)

K562 cells were cross-linked at room temperature for 10 minutes by adding formaldehyde to a final concentration of 1%, followed by quenching with glycine. Cells were washed twice with PBS, pelleted, and stored at -80°C. Pellets were sequentially lysed in LB1 (50 mM HEPES-KOH pH 7.4, 140 mM NaCl, 1 mM EDTA, 0.5 mM EGTA, 10% glycerol, 0.5% NP-40, 0.25% Triton X-100, protease inhibitors), LB2 (10 mM Tris-HCl pH 8.0, 200 mM NaCl, 1 mM EDTA, 0.5 mM EGTA, protease inhibitors), and LB3 (10 mM Tris-HCl pH 8.0, 200 mM NaCl, 1 mM EDTA, 0.5 mM EGTA, 0.1% NaDOC, 0.1% SDS, protease inhibitors). Chromatin was sonicated (Covaris: 5% duty, 200 cycles, 140 PIP, 20 min) to obtain DNA fragments of 100-300 bp. ChIP was performed using an antibody against KAP1. Beads were coated with the antibody at 4°C, followed by overnight incubation with chromatin at 4°C. Immunoprecipitated complexes were washed sequentially with Low Salt Wash Buffer (10 mM Tris-HCl pH 8.0, 1 mM EDTA, 150 mM NaCl, 0.15% SDS), High Salt Wash Buffer (10 mM Tris-HCl pH 8.0, 1 mM EDTA, 500 mM NaCl, 0.15% SDS), LiCl Buffer (10 mM Tris-HCl pH 8.0, 1 mM EDTA, 0.5 mM EGTA, 250 mM LiCl, 1% NP-40, 1% NaDOC), and TE buffer. DNA was eluted and purified using QIAGEN columns. ChIP and input DNA were analyzed by qPCR using specific primers. Enrichment was calculated as a percentage of input or relative to a control region using the ΔΔCt method. Experiments were performed in biological duplicates.

### Chemical perturbation with Hydroxyurea (HU)

Cells were cultured in presence of Hydroxyurea (HU-5mM), for 24h before analysis and release in fresh medium for subsequent analysis.

### Cell cycle analyses

Cell cycle distribution was analyzed by flow cytometry measurement of cellular DNA content using PI staining. Three million cells were collected, washed, and resuspended in 1 volume of ice-cold 1x Dulbecco’s Phosphate Buffer Saline (PBS, BioConcept) before being fixed by the addition of 2.5 volumes of ice-cold 100% ethanol during slow vortexing. After 45 min of incubation at 4°C, cells were washed again in ice-cold PBS and resuspended in 1.5 volume of ice-cold PI staining solution (0.1% triton, 200 μg/ml RNAse A, and 50 μg/ml PI). After 30 min of incubation at room temperature in the dark, the samples were analyzed on a Beckman Coulter Gallios flow cytometer, using Kaluza Analysis software, and quantified using FlowJo single-cell analysis software (FlowJo, LLC).

### Data analysis

Unless otherwise specified, graphs were obtained by Excel software or ggplot2 R package.

### Data Availability

The accession number for the RNA-seq and CUT&Tag data generated in this article is GEO: GSE290774

### Statistical analysis

All statistical tests and numbers of biological replicates are listed in the figure legends. All statistical tests were performed with R.

## Acknowledgments

This work was supported by grants from the European Research Council (KRABnKAP, No. 268721; Transpos-X, No. 694658), the Swiss National Science Foundation (310030_152879 and 310030B_173337), and the Aclon Foundation to D.T. We thank our colleagues from the Tronolab for their inputs throughout the project, and Séverine Reynard for administrative assistance.

## Author Contributions

R.F., and D.T. conceived and planned the study with substantial contribution of F.M., and P.T. R.F., C.R., S.O. and O.R. conducted the experiments. R.F., C.P., E.P., and J.D., performed. the bioinformatic analysis. R.F., C.P and D.T wrote the manuscript.

## Supplemental Information

**Figure S1: Related to Figure 1.**
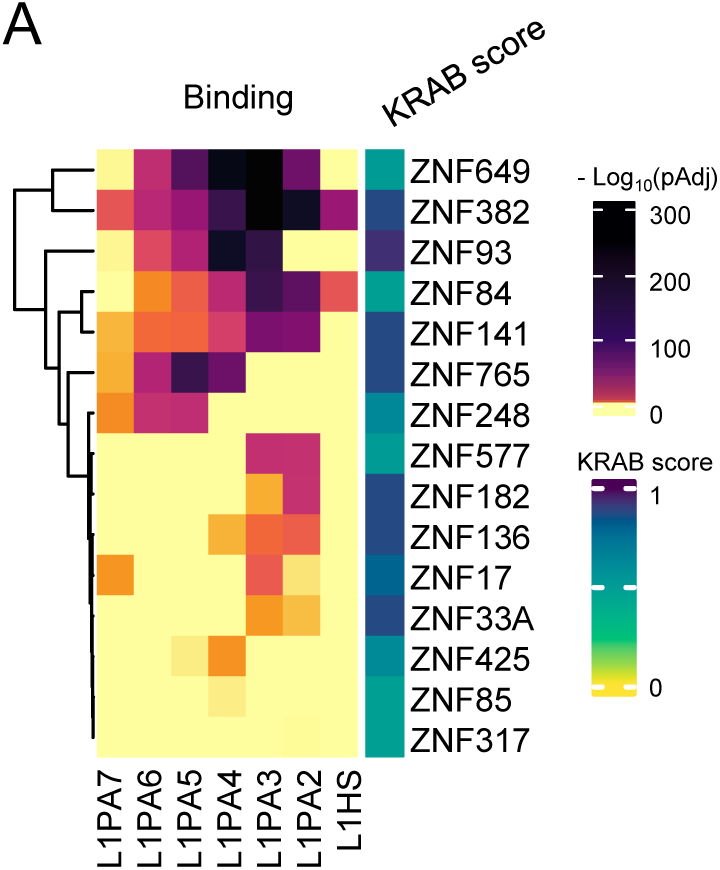
A. Same as Figure 1B, but detailing the seven FLYL1 subfamilies. Rows were clustered using complete-linkage clustering, based on Euclidean distances.

**Figure S2: Related to Figure 2.**
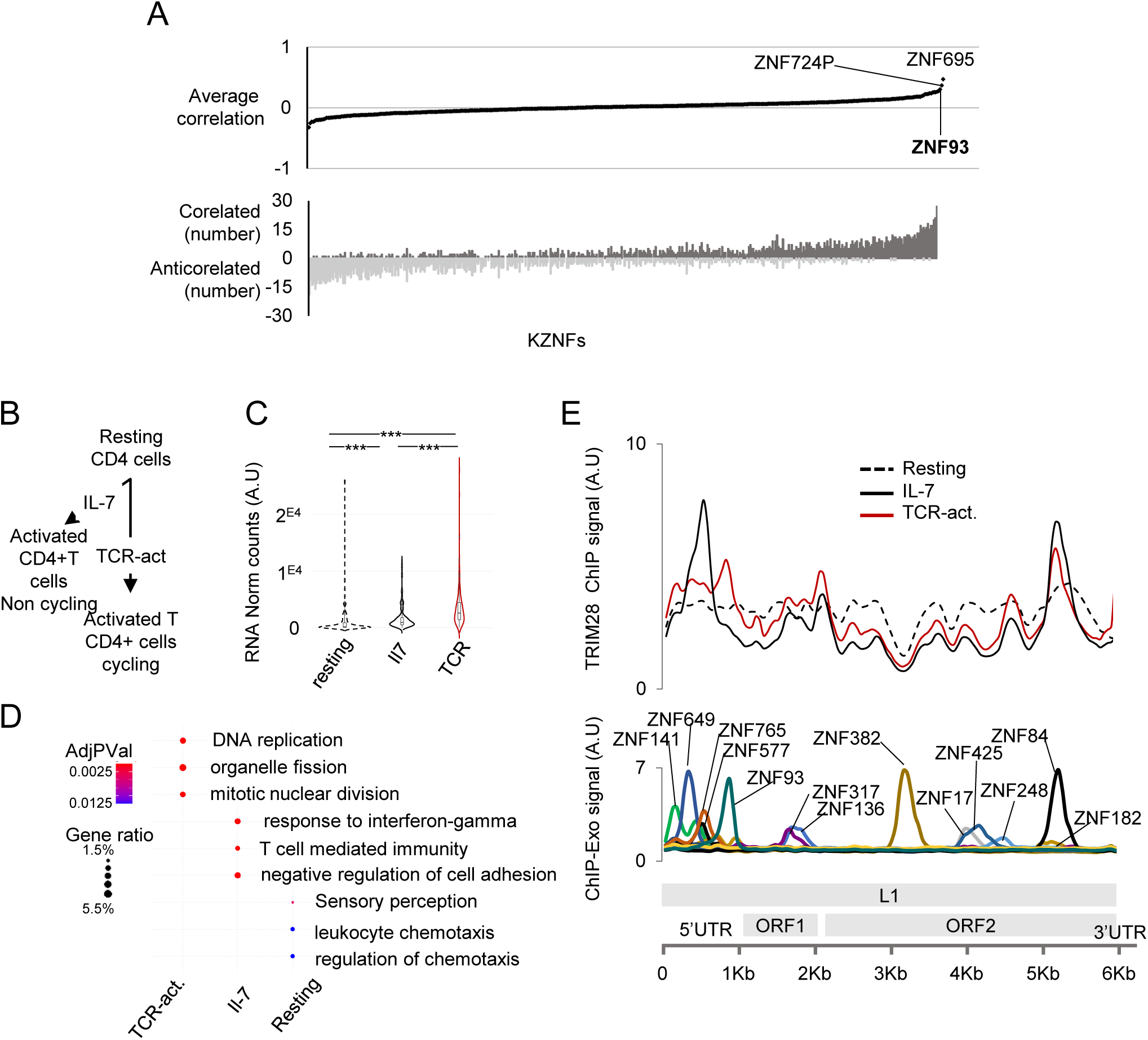
A. Related to Figure 2A. Top: Median correlation of *KZFP* expression with a proliferation signature. Bottom: Number of cancer subtypes where *KZFP* expression is correlated (dark grey) or anticorrelated (light grey) with the proliferation signature. B. Primary resting CD4^+^T cells were activated by T-cell receptor (TCR) crosslinking with a CD3/IL-2 cocktail, leading to activation and proliferation, or treated with IL-7, leading to activation with limited proliferation. C. Normalized counts of proliferation gene signature in resting, IL-7 activated, and TCR-activated CD4^+^T cells. Statistical test: two-tailed t-test. D. Enriched Gene Ontology Biological Processes (GOBP) gene sets in resting, IL-7 activated, and TCR-activated CD4^+^T cells. Dot size represents the ratio of differentially expressed genes in over the total number of genes in each gene set (gene ratio). Dot colour represents the False Discovery Rate (hypergeometric test, FDR < 0.05). E. ChIP signal for TRIM28 across the meta-FLYL1^EN+^ sequence. Figure 1C is reproduced below for comparison.

**Figure S3: Related to Figure 3.**
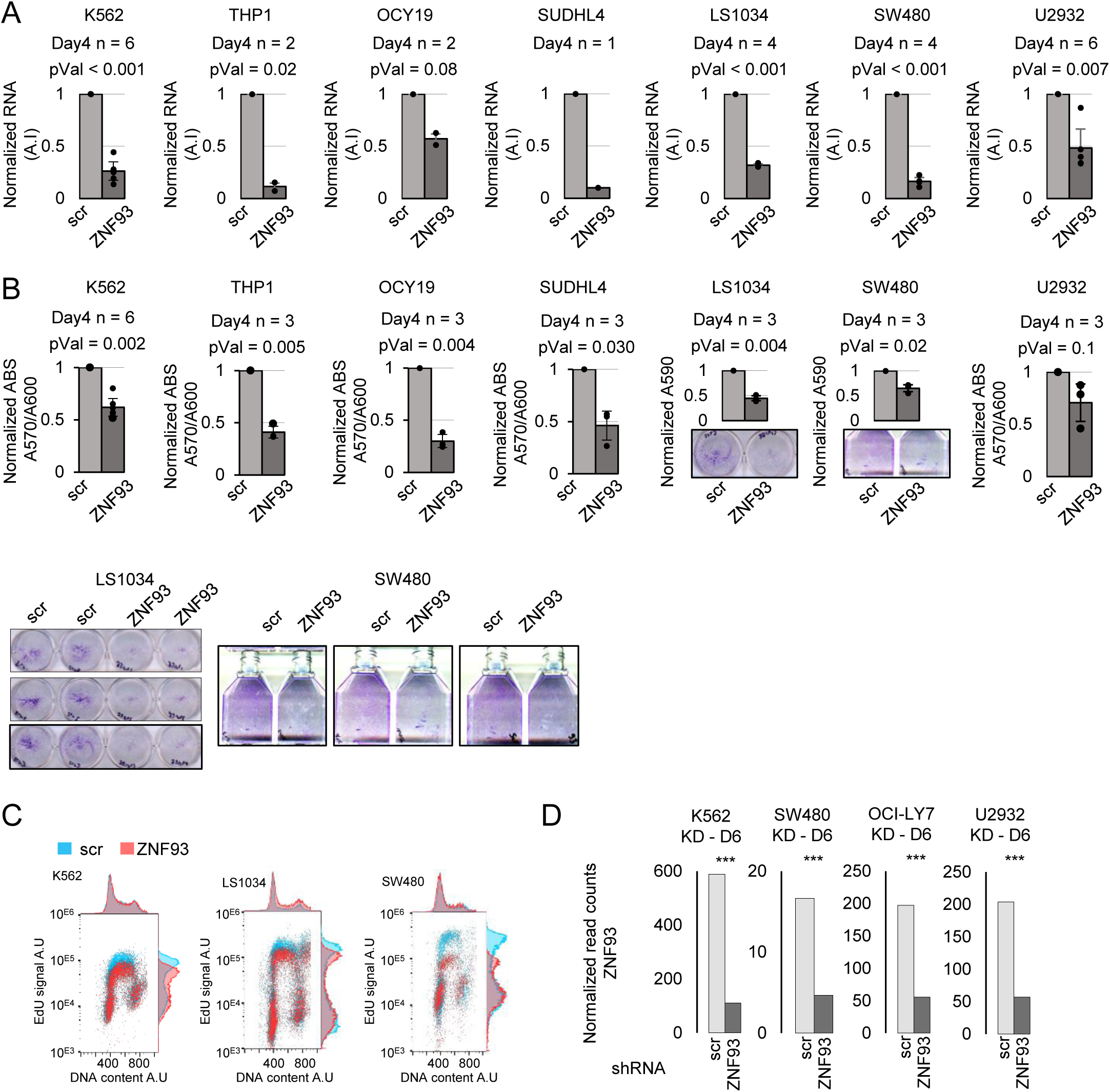
A. Details of Figure 3A: RT-qPCR analysis of *ZNF93* expression six days after *ZNF93* KD in leukemia (K562, THP1), B-cell lymphoma (OCIY19, SUDHL4, and U2932), and colorectal cancer cell lines (LS1034 and SW480). B. Proliferation of the same seven cell lines upon *ZNF93* KD. C. Representative flow cytometry analysis of the replication signal (EdU incorporation intensity, y-axis) and DNA content (DAPI staining, x-axis) upon *ZNF93* KD in K562, SW480, and LS1034 cells. D. ZNF93 depletion efficiency measured by RNA-seq.

**Figure S4: Related to Figure 4.**
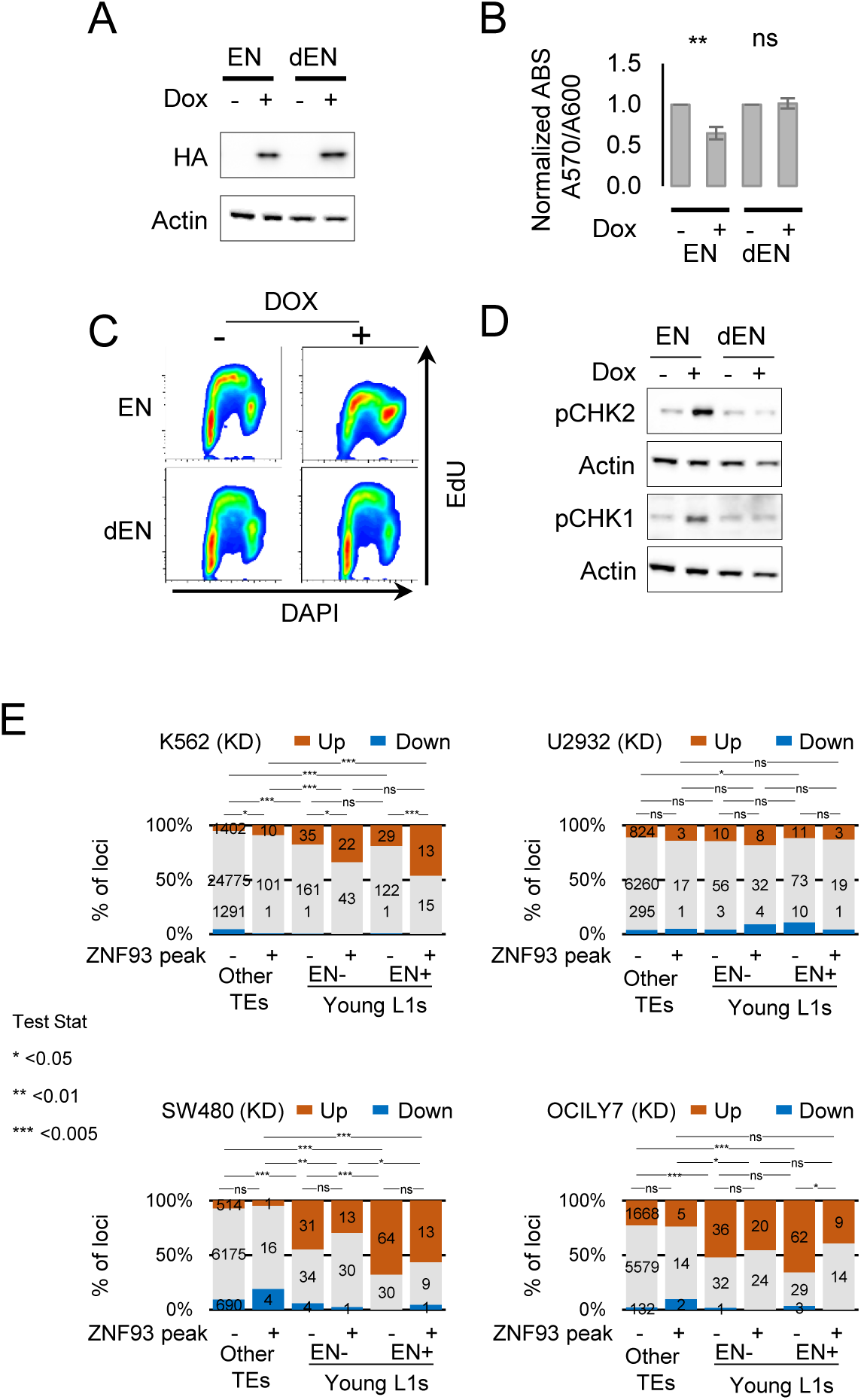
A. HA signal 72 hours after doxycyclin-induced OE of the short version of the ORF2p endonuclease tagged with HA (EN^short^-HA) or its catalytically dead version (EN^short-d^-HA). B. Proliferation assays of K562 cells 96 hours after induction of EN^short^-HA and EN^short-d^-HA (n = 3. two-tailed t-test). C. Flow cytometry analysis of the replication signal (EdU incorporation intensity, y-axis) and DNA content (DAPI staining, x-axis) 72 hours after induction of EN^short^-HA and EN^short-d^-HA in K562 cells. D. pCHEK2 and pCHEK1 signal 72 hours after induction of EN^short^-HA and EN^short-d^-HA. E. Percentage of TEs by differential expression status upon *ZNF93* KD in K562, U2932, OCI-LY7, and SW480. Orange: upregulated; Blue: downregulated (adj. p < 0.05). TEs are categorized into FLYL1^EN+^, FLYL1^EN−^, and other TEs, subcategorized based overlap with ZNF93 peaks (Imbeault et al., 2017). Statistical test: Fischer’s exact test.

**Figure S5: Related to Figure 5.**
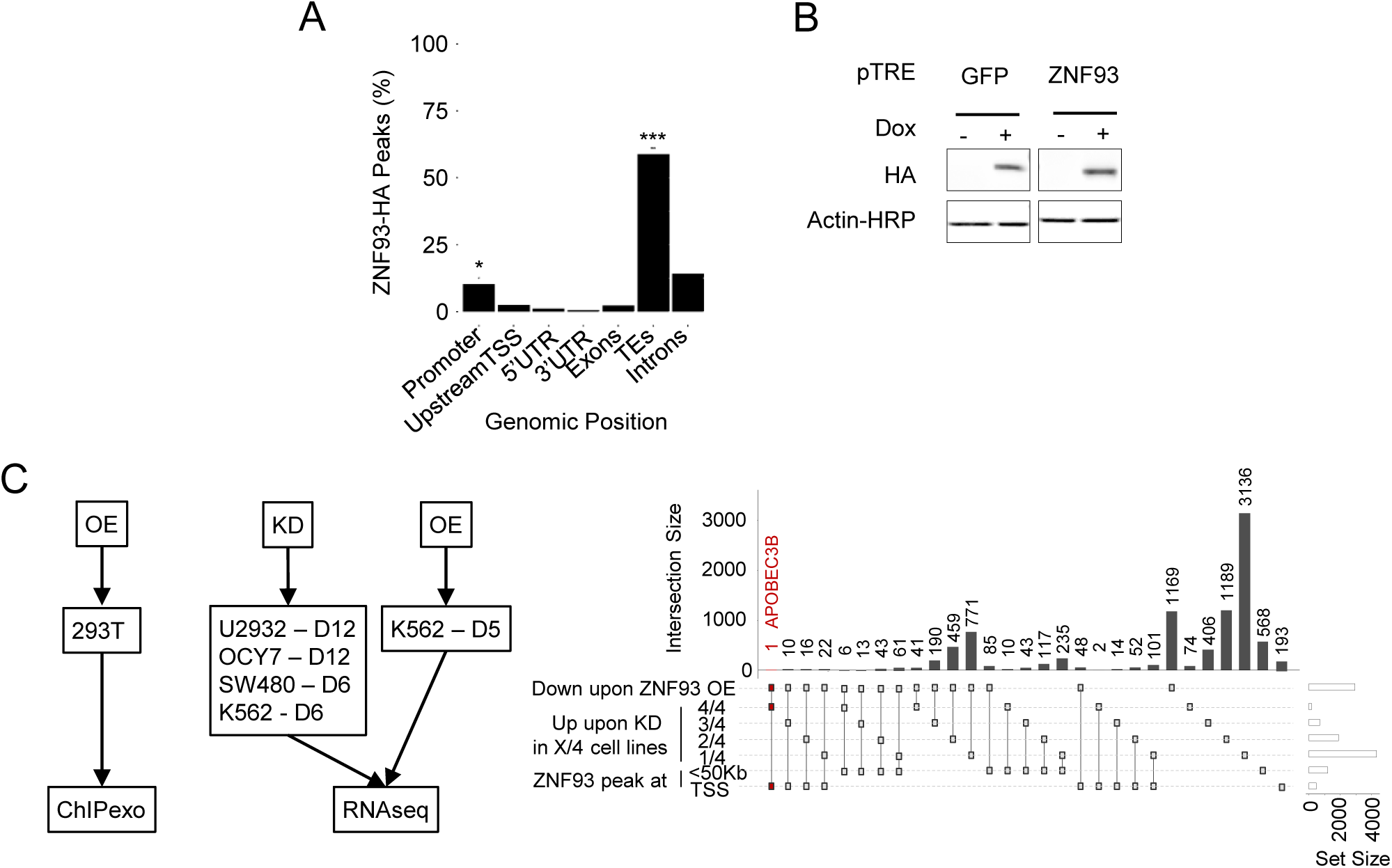
A. % of peaks detected in HEK293T overexpressing ZNF93-HA overlapping with genomic regions. Data from Imbeault *et al.*, 2017. Statistical test: hypergeometric test. B. HA signal after OE of HA-tagged ZNF93 or GFP in K562 cells. C. Left: Datasets used for characterizing ZNF93-dependent transcriptional changes. Right: Intersection of differentially expressed (DE) genes in datasets described in Fig. 5A.

**Figure S6: Related to Figure 6.**
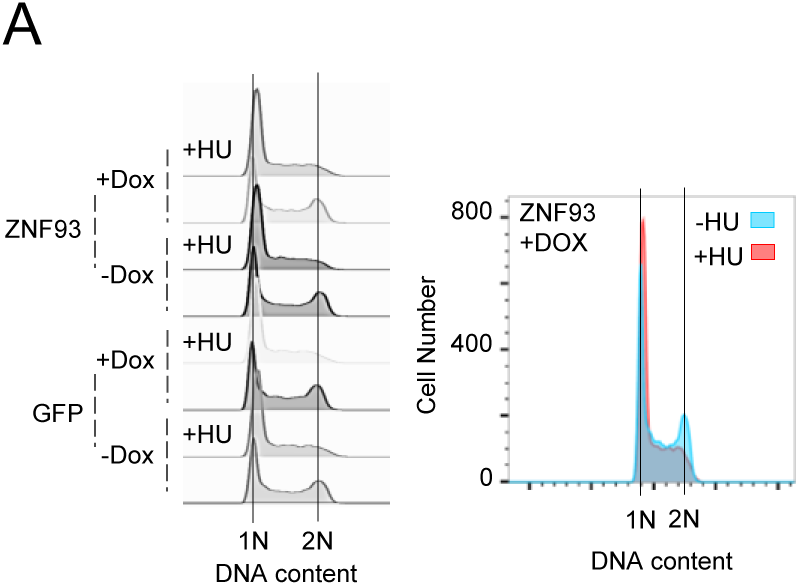
A. Representative flow cytometry analysis of DNA content (propidium iodide incorporation intensity, x-axis) in K562 cells after ZNF93 or GFP overexpression treated with HU.

## REFERENCES

1. A. P. J. de Koning, W. Gu, T. A. Castoe, M. A. Batzer, D. D. Pollock, Repetitive Elements May Comprise Over Two-Thirds of the Human Genome. PLOS Genet. 7, e1002384 (2011).

2. S. Boissinot, A. Sookdeo, The Evolution of LINE-1 in Vertebrates. Genome Biol. Evol. 8, 3485– 3507 (2016).

3. C. Mendez-Dorantes, K. H. Burns, LINE-1 retrotransposition and its deregulation in cancers: implications for therapeutic opportunities. Genes Dev. 37, 948–967 (2023).

4. S. L. Gasior, T. P. Wakeman, B. Xu, P. L. Deininger, The human LINE-1 retrotransposon creates DNA double-strand breaks. J. Mol. Biol. 357, 1383–1393 (2006).

5. J. N. Wells, C. Feschotte, A Field Guide to Eukaryotic Transposable Elements. Annu. Rev. Genet. 54, 539–561 (2020).

6. F. M. J. Jacobs, et al., An evolutionary arms race between KRAB zinc-finger genes ZNF91/93 and SVA/L1 retrotransposons. Nature 516, 242–245 (2014).

7. J. de Tribolet-Hardy, et al., Genetic features and genomic targets of human KRAB-zinc finger proteins. Genome Res. 33, 1409–1423 (2023).

8. O. Rosspopoff, D. Trono, Take a walk on the KRAB side. Trends Genet. TIG 39, 844–857 (2023).

9. M. Imbeault, P.-Y. Helleboid, D. Trono, KRAB zinc-finger proteins contribute to the evolution of gene regulatory networks. Nature 543, 550–554 (2017).

10. P.-Y. Helleboid, et al., The interactome of KRAB zinc finger proteins reveals the evolutionary history of their functional diversification. EMBO J. 38, e101220 (2019).

11. J. Tycko, et al., High-Throughput Discovery and Characterization of Human Transcriptional Effectors. Cell 183, 2020–2035.e16 (2020).

12. B. Brouha, et al., Hot L1s account for the bulk of retrotransposition in the human population. Proc. Natl. Acad. Sci. U. S. A. 100, 5280–5285 (2003).

13. K. B. Chiappinelli, et al., Inhibiting DNA Methylation Causes an Interferon Response in Cancer via dsRNA Including Endogenous Retroviruses. Cell 162, 974–986 (2015).

14. D. Roulois, et al., DNA-Demethylating Agents Target Colorectal Cancer Cells by Inducing Viral Mimicry by Endogenous Transcripts. Cell 162, 961–973 (2015).

15. T. L. Cuellar, et al., ---Silencing of retrotransposons by SETDB1 inhibits the interferon response in acute myeloid leukemia--. J. Cell Biol. 216, 3535–3549 (2017).

16. H. Tunbak, et al., The HUSH complex is a gatekeeper of type I interferon through epigenetic regulation of LINE-1s. Nat. Commun. 11, 5387 (2020).

17. M. De Cecco, et al., L1 drives IFN in senescent cells and promotes age-associated inflammation. Nature 566, 73–78 (2019).

18. J. Z. Shen, et al., FBXO44 promotes DNA replication-coupled repetitive element silencing in cancer cells. Cell 184, 352–369.e23 (2021).

19. D. Ardeljan, et al., Cell fitness screens reveal a conflict between LINE-1 retrotransposition and DNA replication. Nat. Struct. Mol. Biol. 27, 168–178 (2020).

20. P. Mita, et al., BRCA1 and S phase DNA repair pathways restrict LINE-1 retrotransposition in human cells. Nat. Struct. Mol. Biol. 27, 179–191 (2020).

21. S. Wissing, M. Montano, J. L. Garcia-Perez, J. V. Moran, W. C. Greene, Endogenous APOBEC3B restricts LINE-1 retrotransposition in transformed cells and human embryonic stem cells. J. Biol. Chem. 286, 36427–36437 (2011).

22. M. D. Stenglein, R. S. Harris, APOBEC3B and APOBEC3F inhibit L1 retrotransposition by a DNA deamination-independent mechanism. J. Biol. Chem. 281, 16837–16841 (2006).

23. H. P. Bogerd, et al., Cellular inhibitors of long interspersed element 1 and Alu retrotransposition. Proc. Natl. Acad. Sci. U. S. A. 103, 8780–8785 (2006).

24. H. Muckenfuss, et al., APOBEC3 proteins inhibit human LINE-1 retrotransposition. J. Biol. Chem. 281, 22161–22172 (2006).

25. C. J. Warren, J. A. Westrich, K. V. Doorslaer, D. Pyeon, Roles of APOBEC3A and APOBEC3B in Human Papillomavirus Infection and Disease Progression. Viruses 9, 233 (2017).

26. D. W. Cescon, B. Haibe-Kains, DNA replication stress: a source of APOBEC3B expression in breast cancer. Genome Biol. 17, 202 (2016).

27. J. Nikkilä, et al., Elevated APOBEC3B expression drives a kataegic-like mutation signature and replication stress-related therapeutic vulnerabilities in p53-defective cells. Br. J. Cancer 117, 113–123 (2017).

28. L. Xing, et al., APOBEC3B Is Induced By Activation of DNA Repair Pathway and Modulates the Survival and Treatment Response in Human Multiple Myeloma. Blood 132, 407 (2018).

29. S. B. Bader, et al., Replication catastrophe induced by cyclic hypoxia leads to increased APOBEC3B activity. Nucleic Acids Res. 49, 7492–7506 (2021).

30. S. Venkatesan, et al., Induction of APOBEC3 Exacerbates DNA Replication Stress and Chromosomal Instability in Early Breast and Lung Cancer Evolution. Cancer Discov. 11, 2456– 2473 (2021).

31. N. Kanu, et al., DNA replication stress mediates APOBEC3 family mutagenesis in breast cancer. Genome Biol. 17, 185 (2016).

32. L. Wong, A. Sami, L. Chelico, Competition for DNA binding between the genome protector replication protein A and the genome modifying APOBEC3 single-stranded DNA deaminases. Nucleic Acids Res. 50, 12039–12057 (2022).

33. C. Zong, et al., APOBEC3B coordinates R-loop to promote replication stress and sensitize cancer cells to ATR/Chk1 inhibitors. Cell Death Dis. 14, 1–12 (2023).

34. M. Petljak, et al., Mechanisms of APOBEC3 mutagenesis in human cancer cells. Nature 607, 799–807 (2022).

35. K. Butler, A. R. Banday, APOBEC3-mediated mutagenesis in cancer: causes, clinical significance and therapeutic potential. J. Hematol. Oncol.J Hematol Oncol 16, 31 (2023).

36. M. B. Burns, N. A. Temiz, R. S. Harris, Evidence for APOBEC3B mutagenesis in multiple human cancers. Nat. Genet. 45, 977–983 (2013).

37. D. R. Caswell, et al., The role of APOBEC3B in lung tumor evolution and targeted cancer therapy resistance. Nat. Genet. 56, 60–73 (2024).

38. S. Venkatesan, et al., Induction of APOBEC3 Exacerbates DNA Replication Stress and Chromosomal Instability in Early Breast and Lung Cancer Evolution. Cancer Discov. 11, 2456– 2473 (2021).

39. S. Venkatesan, et al., APOBEC3 as a driver of genetic intratumor heterogeneity. Mol. Cell. Oncol. 10, 2014734 (2023).

40. C. Pulver, et al., Evolutionarily recent transcription factors partake in human cell cycle regulation. [Preprint] (2024). Available at: https://www.biorxiv.org/content/10.1101/2024.11.04.621792v1 [Accessed 31 January 2025].

41. F. Marzetta, et al., The KZFP/KAP1 system controls transposable elements-embedded regulatory sequences in adult T cells. [Preprint] (2019). Available at: https://www.biorxiv.org/content/10.1101/523597v1 [Accessed 31 January 2025].

42. D. Ardeljan, et al., LINE-1 ORF2p expression is nearly imperceptible in human cancers. Mob. DNA 11, 1 (2019).

43. M. I. Nielsen, et al., Targeted detection of endogenous LINE-1 proteins and ORF2p interactions. [Preprint] (2024). Available at: https://www.biorxiv.org/content/10.1101/2024.11.20.624490v1 [Accessed 31 January 2025].

44. G. A. Stoll, N. Pandiloski, C. H. Douse, Y. Modis, Structure and functional mapping of the KRAB-KAP1 repressor complex. EMBO J. 41, e111179 (2022).

45. J. L. McCann, et al., APOBEC3B regulates R-loops and promotes transcription-associated mutagenesis in cancer. Nat. Genet. 55, 1721–1734 (2023).

46. F. Coquel, et al., SAMHD1 acts at stalled replication forks to prevent interferon induction. Nature 557, 57–61 (2018).

47. F. Coquel, C. Neumayer, Y.-L. Lin, P. Pasero, SAMHD1 and the innate immune response to cytosolic DNA during DNA replication. Curr. Opin. Immunol. 56, 24–30 (2019).

48. H. Técher, P. Pasero, The Replication Stress Response on a Narrow Path Between Genomic Instability and Inflammation. Front. Cell Dev. Biol. 9, 1671 (2021).

49. A. B. Osmanski, et al., Insights into mammalian TE diversity through the curation of 248 genome assemblies. Science 380, eabn1430 (2023).

50. C. Swanton, N. McGranahan, G. J. Starrett, R. S. Harris, APOBEC Enzymes: Mutagenic Fuel for Cancer Evolution and Heterogeneity. Cancer Discov. 5, 704–712 (2015).

51. P. A. Roelofs, J. W. M. Martens, R. S. Harris, P. N. Span, Clinical Implications of APOBEC3-Mediated Mutagenesis in Breast Cancer. Clin. Cancer Res. Off. J. Am. Assoc. Cancer Res. 29, 1658–1669 (2023).

52. R. G. Vile, A. Melcher, H. Pandha, K. J. Harrington, J. S. Pulido, APOBEC and Cancer Viroimmunotherapy: Thinking the Unthinkable. Clin. Cancer Res. Off. J. Am. Assoc. Cancer Res. 27, 3280–3290 (2021).

53. S. Revathidevi, A. K. Murugan, H. Nakaoka, I. Inoue, A. K. Munirajan, APOBEC: A molecular driver in cervical cancer pathogenesis. Cancer Lett. 496, 104–116 (2021).

54. Z. Duan, et al., ZNF93 increases resistance to ET-743 (Trabectedin; Yondelis) and PM00104 (Zalypsis) in human cancer cell lines. PloS One 4, e6967 (2009).

55. G. Farmiloe, et al., Structural Evolution of Gene Promoters Driven by Primate-Specific KRAB Zinc Finger Proteins. Genome Biol. Evol. 15, evad184 (2023).

56. P. Rice, I. Longden, A. Bleasby, EMBOSS: The European Molecular Biology Open Software Suite. Trends Genet. 16, 276–277 (2000).

57. F. Ramírez, et al., deepTools2: a next generation web server for deep-sequencing data analysis. Nucleic Acids Res. 44, W160–165 (2016).

58. J. De Tribolet-Hardy, et al., Genetic features and genomic targets of human KRAB-Zinc Finger Proteins. [Preprint] (2023). Available at: http://biorxiv.org/lookup/doi/10.1101/2023.02.27.530095 [Accessed 26 November 2024].

59. F. Martins, et al., A Cluster of Evolutionarily Recent KRAB Zinc Finger Proteins Protects Cancer Cells from Replicative Stress–Induced Inflammation. Cancer Res. 84, 808–826 (2024).

60. B. Langmead, S. L. Salzberg, Fast gapped-read alignment with Bowtie 2. Nat. Methods 9, 357– 359 (2012).

61. A. R. Quinlan, I. M. Hall, BEDTools: a flexible suite of utilities for comparing genomic features. Bioinformatics 26, 841–842 (2010).

62. A. Subramanian, et al., Gene set enrichment analysis: A knowledge-based approach for interpreting genome-wide expression profiles. Proc. Natl. Acad. Sci. 102, 15545–15550 (2005).

63. V. K. Mootha, et al., PGC-1α-responsive genes involved in oxidative phosphorylation are coordinately downregulated in human diabetes. Nat. Genet. 34, 267–273 (2003).

